# Reactive astrocytes associated with prion disease impair the blood brain barrier

**DOI:** 10.1101/2023.03.21.533684

**Authors:** Rajesh Kushwaha, Yue Li, Natallia Makarava, Narayan P. Pandit, Kara Molesworth, Konstantin G. Birukov, Ilia V. Baskakov

**Author notes:** To whom correspondence should be addressed: Center for Biomedical Engineering and Technology, University of Maryland School of Medicine, 111 S. Penn St., Baltimore, MD 21201.

## Abstract

**Background:** Impairment of the blood-brain barrier (BBB) is considered to be a common feature among neurodegenerative diseases, including Alzheimer’s, Parkinson’s and prion diseases. In prion disease, increased BBB permeability was reported 40 years ago, yet the mechanisms behind the loss of BBB integrity have never been explored. Recently, we showed that reactive astrocytes associated with prion diseases are neurotoxic. The current work examines the potential link between astrocyte reactivity and BBB breakdown.

**Results:** In prion-infected mice, the loss of BBB integrity and aberrant localization of aquaporin 4 (AQP4), a sign of retraction of astrocytic endfeet from blood vessels, were noticeable prior to disease onset. Gaps in cell-to-cell junctions along blood vessels, together with downregulation of Occludin, Claudin-5 and VE-cadherin, which constitute tight and adherens junctions, suggested that loss of BBB integrity is linked with degeneration of vascular endothelial cells. In contrast to cells isolated from non-infected adult mice, endothelial cells originating from prion-infected mice displayed disease-associated changes, including lower levels of Occludin, Claudin-5 and VE-cadherin expression, impaired tight and adherens junctions, and reduced trans-endothelial electrical resistance (TEER). Endothelial cells isolated from non-infected mice, when co-cultured with reactive astrocytes isolated from prion-infected animals or treated with media conditioned by the reactive astrocytes, developed the disease-associated phenotype observed in the endothelial cells from prion-infected mice. Reactive astrocytes were found to produce high levels of secreted IL-6, and treatment of endothelial monolayers originating from non-infected animals with recombinant IL-6 alone reduced their TEER. Remarkably, treatment with extracellular vesicles produced by normal astrocytes partially reversed the disease phenotype of endothelial cells isolated from prion-infected animals.

**Conclusions:** To our knowledge, the current work is the first to illustrate early BBB breakdown in prion disease and to document that reactive astrocytes associated with prion disease are detrimental to BBB integrity. Moreover, our findings suggest that the harmful effects are linked to proinflammatory factors secreted by reactive astrocytes.

## Background

Chronic neuroinflammation is recognized as one of the main characteristics of neurodegenerative diseases, including Alzheimer’s, Parkinson’s and prion diseases, and involves the sustained transformation of homeostatic states of microglia and astrocytes into reactive states [1]. A number of reactive phenotypes of glia, ranging from largely protective to highly toxic with respect to their effect on neuronal survival, have been described in different neurodegenerative diseases, disease stages, and animal models [2–7]. However, regardless of whether the net effects of reactive microglia or astrocytes are positive or negative, neuroinflammation is often associated with a breakdown of the blood-brain barrier (BBB), which is also considered to be a common feature among neurodegenerative diseases (reviewed in [8]). The BBB consists of endothelial cells within brain microvessels with tightly sealed cell-to-cell junctions. The integrity of the BBB is essential for brain homeostasis, as it protects the CNS from the entry of pathogens and plasma factors that could be toxic to neurons. Compromised BBB integrity is harmful due to increased transmigration of leukocytes and macrophages, invasion of pathogens, and entry of blood-derived molecules [8].

Prion diseases, also known as transmissible spongiform encephalopathies, are lethal and transmissible neurodegenerative disorders that affect both humans and animals [9]. Prions, or PrP^Sc^, are proteinaceous infectious agents that consist of misfolded, self-replicating states of a sialoglycoprotein called the prion protein or PrP^C^ [9, 10]. Prions or PrP^Sc^ are transmitted between organisms or from cell to cell by recruiting host-encoded PrP^C^ and replicating their misfolded structures via a template-assisted mechanism [11]. PrP^C^ is a sialoglycoprotein that is posttranslationally modified with up to two N-linked sialoglycans and a GPI anchor [12–17]. Although prion strains consist of the same protein, they differ in their structure and composition of sialoglycoforms [18–21]. Prion strains induce multiple disease phenotypes characterized by different, strain-specific PrP^Sc^ deposition patterns, incubation times to disease and brain areas affected by prions [22, 23]. Regardless of strain-specific structures or disease phenotype, chronic neuroinflammation is regarded as a common feature of the disease [24].

Increased BBB permeability associated with prion diseases was noticed nearly 40 years ago in prion-infected mice, and was found to be common among animals infected with different prion strains [25]. Significant caspase immunoreactivity of blood vessels, indicative of endothelial cell death, was observed in the brains of prion-infected mice [26]. In recent studies, a mouse model with compromised BBB was employed to examine the role of BBB permeability on prion neuroinvasion [27]. Upon infection via peripheral routes, BBB permeability did not dictate the timing of entry of prions to the brain or disease initiation [27]. However, the question of whether and how prion infection of the CNS causes BBB breakdown has never been addressed. In fact, there is a critical gap in our understanding of the underlying causes of BBB breakdown in prion and other neurodegenerative diseases. Unlike most neurodegenerative diseases, authentic prion diseases can be induced in wild-type or inbred animals via transmission of PrP^Sc^ [28].

Under normal conditions, astrocytes play an essential role in the development and maintenance of the BBB [29–32]. However, whether in prion disease, astrocytes continue to support the BBB upon their transformation into a reactive state or are harmful to the BBB has never been explored. [33]. In mice infected with prions, transcriptome analysis revealed significant perturbation in the expression of astrocyte-specific genes involved in BBB maintenance [34]. Endothelial cells of blood vessels are wrapped by astrocytic endfeet that form tight contacts between astrocytes and perivascular basal lamina [35]. Aquaporin 4 (Aqp4) is the most prevalent water channel that localizes on astrocytic endfeet surfaces and is regarded as a key channel in maintaining water homeostasis in the CNS [36]. Recently, we reported changes in the cellular localization of Aqp4 in reactive astrocytes in prion disease [34]. Moreover, accumulation of Aqp4 within or around prion plaques was found in human prion diseases [37]. These changes suggest a loss of astrocyte polarization and possible dysregulation of astrocyte functions responsible for BBB maintenance. Here, we hypothesize that astrocyte reactivity, while originating as a physiological response to an insult, gives rise to a disease-associated phenotype that interferes with astrocyte homeostatic functions, among which is BBB maintenance.

In the current study, we examined the potential link between astrocyte reactivity and BBB breakdown. We observed aberrant AQP4 localization along with an increase in BBB permeability at pre-symptomatic stages of the disease. Endothelial cells isolated from prion-infected animals displayed a disease-associated phenotype characterized by downregulation of proteins that constitute tight and adherens junctions, and a loss of cell-to-cell junctions. Remarkably, reactive astrocytes or media conditioned by reactive astrocytes isolated from prion-infected mice induced a disease-associated phenotype in endothelial cells originating from non-infected adult mice. Vice versa, extracellular vesicles (EVs) produced by normal astrocytes partially reversed the disease-associated phenotype of endothelial cells isolated from prion-infected mice. This study is the first to document that, in prion disease, reactive astrocytes drive pathological changes in BBB.

## Methods

### Reagents and kits

Evans blue dye, Poly-L-lysine (PLL), Poly-D-lysine (PDL), collagenase from clostridium histolyticum, sodium bicarbonate, paraformaldehyde (PFA), bovine serum albumin (BSA), normal goat serum (NGS), tween 20, triton-X-100, protease inhibitor cocktail (PIC), CellLytic MT mammalian cell lysis buffer, ponceau S, dimethyl sulfoxide, FITC-dextran, hydrochloric acid, puromycin and endothelial cell growth supplement were purchased from Sigma Chemical Co. (St. Louis, MO). Trypsin-EDTA, Dulbecco’s modified eagle medium: F12 (DMEM/F12), Hank’s balanced salt solution (HBSS), phosphate buffer saline (PBS), Dulbecco’s phosphate buffered saline (DPBS), antibiotic-antimycotic, penicillin/streptomycin, heparin, fetal bovine serum (FBS), fetal bovine serum-exosome-depleted, glutaMAX and protein ladder were purchased from Invitrogen (Carlsbad, CA). Adult mouse brain dissociation kit, myelin removal solution, LS column and C-tubes were from Miltenyi Biotec (Bergisch Gladbach, Germany). VECTASHIELD mounting medium with DAPI was from Vector Laboratories (Burlingame, CA) and Supersignal West pico PLUS Chemiluminescent Substrate was purchased from Thermo Scientific (Rockford, IL). Aurum Total RNA Mini Kit, SYBR Green and iScript cDNA Synthesis Kit were procured from Bio-Rad laboratories (Hercules, CA, CA). ELISA kit for IL-6 and recombinant mouse IL-6 wrtr from R&D Systems (Minneapolis, MN). Bicinchoninicacid (BCA) protein assay kit, 70 μm nylon mesh filter, 0.22µm filters, MTT (3-(4,5-dimethylthiazol-2-yl)-2,5-diphenyltetrazolium bromide), polyvinylidenefluoride (PVDF) membrane were procured from Millipore (Temecula, CA). Collagenase/Dispase was from Roche (Basel, Switzerland). Mouse collagen type I and collagen type IV from Corning Life Sciences (Durham, NC). ECIS culture ware, 8W10E PET was procured from Applied Biophysics (Troy, NY).

### Antibodies

Rabbit polyclonal antibody to AQP4 (cat. HPA014784) was from Millipore Sigma (Rockville, MD). Mouse monoclonal antibody to VE-cadherin (cat. sc9989) was from Santa Cruz Biotechnology (Dallas, TX). Rabbit polyclonal antibodies to Claudin-5 (cat. 34-1600), Occludin (cat. 71-1500), ZO-1 (cat. 61-7300) and VE-Cadherin (cat. 36-1900) were from Thermo Fisher Scientific (Waltham, MA). Rabbit monoclonal antibody to Flotillin-1 (cat. 3253) and Alix (cat. 2171) were from Cell Signaling Technology (Danvers, MA). Mouse monoclonal antibody to CD31 (cat. ab24590) and olig2 (cat. ab136253) were from Abcam (Cambridge, MA). Chicken polyclonal antibody to GFAP (cat. AB5541), mouse monoclonal antibodies to β-actin (cat. A5441), horseradish peroxidase (HRP) conjugated secondary anti-rabbit IgG (cat. A0545) and anti-mouse IgG (cat. A9044) were procured from Sigma-Aldrich (St. Louis, MO). Rabbit polyclonal antibody to Iba1 (cat. 01919741) was from Wako (Richmond, VA). Alexa Fluor 488 goat anti-rabbit IgG, Alexa Fluor 488 goat anti-chicken IgG, Alexa Fluor 488 goat anti-mouse IgG, Alexa Fluor 546 goat anti-rabbit IgG and Alexa Fluor 546 goat anti-mouse IgG secondary antibodies were purchased from Invitrogen (Carlsbad, CA).

### Animals

C57BL/6J male and female mice were housed in a 12 hour day and light cycle environment with the *ad libitum* availability of diet and water. Six-week-old C57BL/6J male and female mice were intraperitoneally inoculated with 200 µl volume of 1% SSLOW brain homogenate in PBS under anesthesia conditions. For the preparation of SSLOW inoculums, the 6^th^ passage of SSLOW in C57BL/6J mice was used [38]. Animals were regularly observed and scored for disease progression using the following signs: hind-limb clasping, ataxia, kyphosis, abnormal gait and loss of weight. Animals were euthanized at pre-symptomatic (92-110 dpi) and terminal (146-185 dpi) stages of the disease. For primary cultures, 70 SSLOW-infected and 70 aged-matched control mice were used.

### BBB permeability assay

Evans blue (EB) dye was used to assess the blood-brain barrier (BBB) permeability in a mouse brain as described earlier [39, 40]. Briefly, SSLOW-infected, aged-matched control and aged mice were anesthetized by isoflurane, then 3% EB in normal saline was inject slowly through the tail vein (4 ml/kg) and allowed to circulate for 3-4 hours. Prior to brain isolation, mice were transcardially perfused using normal saline to remove EB dye from circulation. Brains were post-fixed in 4% PFA and cryoprotected in 30% sucrose. 10 μm thick cryosections of the cerebral cortex and hippocampus were prepared using a vibratome (Leica VT1200, Wetzlar, Germany). The sections were Hoechst-stained, treated with Vectashield mounting medium, and EB fluorescence was analyzed using an inverted Nikon Eclipse TE2000-U microscope (Nikon Instruments Inc.). For analysis of EB absorbance, the EB-injected mice were transcardially perfused, cerebral cortex and hippocampus were dissected, weighed, incubated in formamide solution and then homogenized. Homogenates were incubated overnight at 37 °C and then centrifuged at 12000□g for 30□min to collect the supernatant. The EB content of supernatants was measured by taking optical density at 620 nm using Tecon Microplate Reader (Männedorf, Switzerland) and expressed as μg of EB per gram of tissue.

### Immunohistochemistry

Aged-matched control and terminally ill SSLOW-inoculated mice were anesthetized and perfused with normal saline. The whole brain was removed, postfixed in formalin overnight, and treated for 1 hour in 96% formic acid, then 10 μm cortical sections were prepared using a vibratome (Leica VT1200, Wetzlar, Germany). The same plane and position of cortical sections were maintained across animal groups. To expose epitopes, slides were subjected to 20 min of hydrated autoclaving at 121°C in trisodium citrate buffer, pH 6.0, with 0.05% Tween 20. Sections were blocked with 5% BSA in PBS, and then probed with anti-CD31 (1:500), claudin-5 (1:500), occludin (1:500), ZO-1 (1:500), ve-cadherin (1:500), GFAP (1:1000) or AQP4 (1:500) primary antibodies overnight at 4□°C. Sections were washed three times in PBST and incubated with Alexa fluor secondary antibodies at double dilutions relative to respective primary antibodies for 2 h, washed and mounted in VECTASHIELD media. Four-to-five cortical sections per individual brain were analyzed with an inverted Nikon Eclipse TE2000-U microscope (Nikon Instruments Inc.) equipped with an X-cite 120 illumination system (EXFO Photonics Solutions Inc.) and a cooled 12-bit CoolSnap HQ CCD camera (Photometrics). Images were processed using ImageJ software (NIH). Integrated fluorescence intensity and length of vessels were analyzed and calculated through ImageJ software.

### Ratio of perivascular to parenchymal AQP4

AQP4 distribution was measured on the images taken with 20x objective. 50 pixels long linear profiles were drawn across cortical microvessels, avoiding cell bodies and positioning the vessel in the middle. The parenchymal AQP4 signal was reported as an average intensity of the first and last 15 pixels of each profile. Maximum intensity of the 20 pixels in the middle of each plot represented the perivascular AQP4 signal. The minimum value of each profile was subtracted as a background value. The individual ratios of perivascular to parenchymal AQP4 signal were plotted, and the statistical significance of the difference between infected and normal brains was estimated by Mann-Whitney test in GraphPad Prizm 9.5.1.

### Primary endothelial cultures

Cultures of primary adult mouse brain endothelial cells (BECs) derived from the SSLOW-prion infected or aged-matched control C57BL/6J mice were prepared as previously described with the following modifications [41]. One brain was used per individual culture preparation. After euthanasia and extraction from the skull cavity, brains were gently transferred to a 60 mm petri dish, rinsed with cold DPBS to remove adhering blood, then the midbrain, cerebellum, and olfactory bulb were removed. After removal of meninges, brain cortices were dissociated and digested with the pre-warmed digestion mixture containing 1 mg/mL collagenase/dispase and 10 μg/mL DNAse I for 60 min at 37°C. Digested tissues were pelleted by centrifugation at 250 g for five minutes at 4°C and the obtained pellet was resuspended in 22% (w/v) bovine serum albumin, then centrifuged at 1250 g for 10 min. Myelin on the top of tubes was carefully removed, then the cell pellets were washed with DMEM and centrifuged at 200 g for 5min. The supernatant was aspirated and the cell pellets were again resuspended in a pre-warmed digestion mixture for 30 min at 37 °C. After digestion, cells were pelleted by centrifugation at 250 g for 5 min and washed with DMEM to remove the traces of enzymes and debris. The resulting cells pellets were resuspended in complete endothelial cell growth media (ECGM: DMEM/f12 containing 365 µg/ml L-glutamine and 1 mM sodium pyruvate; 5% heat inactivated FBS, 100 U/ml penicillin, 100 µg/ml streptomycin, 100 U/ml heparin and endothelial cell growth supplement). Then, cells were seeded on collagen-coated chamber slides or culture flasks at plating density 3-5×10^4^ per well or 7×10^5^ per flask and grown in a humidified CO_2_ incubator at 37°C with 5% CO_2_. To obtain pure endothelial cells, puromycin (4 μg/ml) was added to the culture media between days 1–3 to remove non-endothelial cells. Endothelial cell purity was estimated to be > 95% as determined by CD31 immunostaining (Fig. S2).

For examining the effect of astrocytes conditioned medium (ACM) on endothelial cell viability, the cells were treated with ACMs for 0-72h. For assessing the effects of ACM on endothelial junctional proteins and TEER levels, the endothelial cells were treated with ACMs for 72 hours.

### Primary astrocyte culture

Primary cortical astrocyte cultures were isolated from SSLOW-infected terminal or age-matched control C57BL/6J mice following the published protocol [6]. Briefly, after dissection, brains were rinsed with a cold DPBS to remove adhering blood. After removal of meninges, the brain cortices were dissociated and digested using papain-based enzymes dissociation solution and incubated on the gentleMACS octo system for 30 min, according to manufacturer’s instruction (Miltenyi biotec, Germany). Following digestion, cells were resuspended in a buffer containing DPBS, 100 U/ml penicillin and 100 mg/ml streptomycin, then non-cellular debris was removed by passing the cell suspension through a 70 µm nylon single-cell strainer. Then, the clear suspension was centrifuged for 10 min at 250 g, and the obtained pellet was incubated with myelin removal solution at 4°C for 10 min and centrifuge at 250 g for 5 min. The obtained pellet was re-suspended in complete astrocyte growth media (DMEM/F12 containing 365 µg/ml L-glutamine and 1 mM sodium pyruvate; 5% heat inactivated FBS; 100 U/ml penicillin; 100 µg/ml streptomycin). Cells were seeded onto poly-L-lysine (PLL)-coated chamber slides, Transwell inserts or culture flasks at a plating density of 3-4×10^4^ per well, 1.5× 10^5^ cell/cm^2^ in Transwell insert or 7×10^5^ per flask, and grown in humidified CO_2_ incubator at 37°C with 5% CO_2_. The next day after plating the cells, complete media was replaced to remove debris and unattached dead cells.

### *In-vitro* Blood Brain Barrier model

To construct *in vitro* BBB models, the contact co-culture model was established as previously described [42]. Primary astrocytes isolated from SSLOW-infected or age-matched control mice were seeded at a plating density of 1.5× 10^5^ cell/cm^2^ on the bottom sides of the collagen-coated polyester membranes of the Transwell inserts (Corning Life Sciences, Cambridge, MA) and allowed to attach and proliferate for one day. Then, the inserts were inverted to their original orientation and cultured in astrocyte media. Primary endothelial were seeded at a plating density of 1.5× 10^5^ cell/cm^2^ on the inside side of the Transwell inserts and placed in wells of the 24-well culture plates. The astrocyte-endothelial co-cultures were cultured in ECGM supplemented with 2.5% FBS. The medium was replaced every other day until cells reached 90% confluency. BBB models could be cultured for up to 15 days.

### Preparation of conditioned medium

To prepare astrocyte-conditioned media (ACM), primary astrocytes were plated at a density of 7×10^5^ per culture flask and grown in humidified CO_2_ incubator at 37°C as described above. After achieving 50-60% confluency, the monolayers of astrocytes were subjected to an endothelial growth media to generate ACM. Media was replenished every 2 days. After 70-80% confluency (2-3 weeks), media were collected and centrifuged at 1000 rpm for 5 min to remove cellular debris and used immediately.

### Isolation and purification of extracellular vesicles (EVs)

EVs were isolated from the astrocyte culture medium using ultracentrifugation as previously described [43]. Briefly, astrocytes were cultured in flasks (7×10^5^). The astrocyte culture medium was replaced with the medium containing 10% exosome-free FBS. After 24 h incubation, the astrocyte culture medium was transferred to an ultracentrifuge tube and centrifuged sequentially at 300 g for 10 min, 2000 g for 10 min, 10,000 g for 30 min, and 100,000 g for 70 min (Beckman, SW 40 rotor). The obtained pellets were resuspended in cold PBS and ultracentrifuged at 100,000g for an additional 70 min. The EV-containing pellets were resuspended in PBS.

### Cell viability assay

To assess viability of endothelial cells, MTT (3-(4,5-dimethylthiazol-2-yl)-2,5-diphenyltetrazolium bromide) assay was performed per manufacturer protocol. Briefly, primary endothelial cells were seeded in collagen-coated 96-well culture plates at a density of 1×10^4^ cells per well in triplicates, then grown until 70-80% confluency and were treated with control or SSLOW-astrocytes conditioned medium for up to 72 hours. The culture medium was cautiously aspirated, then MTT was added and incubated at 37°C for 4h in a 5% CO_2_ incubator. After adding 100 μl per well of detergent reagent, the plate was kept for 2 h in the dark at room temperature, then absorbance was measured at 570 nm using a Microplate reader (Tecan, infinite 200 PRO, Switzerland).

### Trans-endothelial Electrical Resistance (TEER)

The barrier properties of endothelial cells were analyzed by measurements of trans-endothelial electrical resistance (TEER) across confluent monolayers using an electrical cell-substrate impedance sensing instrument (Applied Biophysics, Troy, NY) as previously described [44]. Briefly, brain endothelial cells were cultured on small gold microelectrodes (8 well chamber slides, ECIS, Applied biosystem, Waltham, MA) in endothelial growth media containing 5% FBS. Before the experiment, growth media was replaced with serum free media, then a 4,000-Hz AC signal with 1-V amplitude was applied across cell monolayers. After electrical resistance achieved a steady state level of approximately 1000 Ω, endothelial monolayers were analyzed for 24 h. The total electrical resistance was measured dynamically through monolayers and was defined as the combined resistance between the basal surface of the cells and the electrode, reflective of focal adhesion, and the resistance between the cells.

### Immunocytochemistry

Endothelial and astrocyte cells were fixed in 4% paraformaldehyde for 15 min at room temperature, washed with PBS, permeabilization in methanol for 30 min, and washed again with PBS. Cells were blocked with 5% bovine serum albumin for 2 h, then incubated with primary antibodies at the following dilutions: Occludin 1 :400, claudin-5 1:500, VE-cadherin 1:500, CD31 1:1000, Iba1 1:1000, NeuN 1:1000, Olig2 1:500 and GFAP 1:500 overnight at 4°C. Cells were washed with PBST buffer (PBS+0.1% TWEEN-20), then incubated with Alexa Fluor 488 goat anti-mouse IgG, Alexa Fluor 546 goat anti-mouse IgG, Alexa Fluor 488 goat anti-rabbit IgG or Alexa Fluor 546 goat anti-rabbit IgG conjugates for 2 hours and mounted in VECTASHIELD mounting medium with DAPI (Vector Laboratories, Burlingame, CA). Fluorescence images were taken using an inverted Nikon Eclipse TE2000-U microscope. Fluorescence integrated intensity and uninterrupted length of cell-to-cell junctions were analysed using Image-J 1.453t software (http://rsb.info.nih.gov/ij/; Wayne Rasband, National Institutes of Health, Bethesda, MD).

### Analysis of astrocytes morphology

For identifying the area, perimeter and number of processes in astrocytes, images of non-overlapping GFAP-positive astrocytes were taken using an inverted Nikon Eclipse TE2000-U microscope and analysed using ImageJ software. Images of 5 random fields of view were selected per well of chamber slides; 15 to 20 images per chamber slide per one experimental condition were taken. After background subtraction and threshold adjusting, 5-6 non-overlapping cells per field of view were analyzed.

### RT-qPCR

Total RNA was isolated from primary endothelial cells, astrocytes or brain cortical tissue using Aurum Total RNA Mini Kit (Bio Rad, Hercules, CA) according to the manufacturer’s instruction. Total RNA was dissolved in the elution buffer, then the quantity and purity of mRNA were analyzed using NanoDrop ND-1000 Spectrophotometer (Thermo Fisher Scientific, Waltham, MA). Complementary DNA (cDNA) synthesis was performed using iScript cDNA Synthesis Kit as described elsewhere. The cDNA was amplified with CFX96 Touch Real-Time PCR Detection System (Bio-Rad, Hercules, CA) using SsoAdvanced Universal SYBR Green Supermix and primers listed in Table S1. The PCR protocol consist of 95°C for 2 min followed by 40 amplification cycles at 95°C for 5 s and 60°C for 30 s. Optimum primer pairs and internal housekeeping control gene, glyceraldehydes 3-phosphate dehydrogenase (*GAPDH*) was designed using Primer Express version 2.0.0 (Table 1). The data were analyzed using CFX96 Touch Real-Time PCR Detection System Software. The ΔCt for each RNA sample was calculated by subtracting the mean Ct of the housekeeping gene, *GAPDH,* from the mean Ct of the gene of interest and then relative mRNA gene expression was calculated using 2^-ΔΔCt^ method.

### Enzyme-linked immunosorbent assay (ELISA)

The IL-6 levels in astrocytes conditioned medium (ACM) were determined using the IL-6 ELISA kit per the manufacturer’s instruction (R&D Systems, Minneapolis, MN). Briefly, primary astrocytes were plated at a density of 7×10^5^ per culture flask, grown in a humidified CO_2_ incubator at 37°C, and conditioned media collected as described above.

### Protein extraction and Western blotting

Primary endothelial and astrocytes cells and cortical tissue samples were homogenized using a lysis buffer containing a protease inhibitor cocktail. The lysates were centrifuged (4°C, 20000 g) for 30 min. Protein concentration was determined using BCA assay per the manufacturer’s instructions. Protein samples were prepared with 1X SDS sample loading buffer and denatured at 85°C for 15 min. An equal amount of protein samples (40µg) along with a pre-stained protein ladder (Marker) were loaded onto 10%-12% tris-glycine polyacrylamide gel and run in 1X running buffer at 100V. After completion of electrophoresis, the transfer onto the PVDF membrane (activated in methanol) was conducted at 16 V for 60 min. Then membranes were washed with TBST (10mM Tris, pH 8.0, 150 mM NaCl and 0.01% Tween 20), blocked with 5% non-fat milk for 1 h, washed thrice with TBST and probed overnight with CD31 (1:3000), Claudin-5 (1:2000), Occludin (1:2000), ZO-1 (1:3000), VE-Cadherin (1:2000), Flotillin-1 (1:3000), Alix (1:3000), GFAP (1:3000) and β-actin (1:10,000) antibodies. Then, membranes were washed four times with TBST and incubated with HRP-conjugated secondary antibodies at double dilutions relative to the dilutions of primary antibodies. Protein bands were visualized using Supersignal West pico Maximum Sensitivity Substrate and FlourChem M imaging system (Protein Simple). Densitometry analysis was performed using Bio-Rad Quantity One image analysis software (Bio-Rad, Hercules, CA).

### FITC-Dextran permeability assay

Permeability assays of endothelial-astrocyte co-cultures were performed in 24-well culture plates with transwell inserts. Upon reaching 70-80% confluence, inserts were equilibrated in the assay medium (phenol red-free DMEM supplemented with 1% FBS) for 30 min at 37°C. 10 kDa FITC-conjugated dextran (5mg/mL, Sigma-Aldrich) was applied to the luminal compartments, then 100 µl aliquots were collected from the abluminal compartment after 2 and 4 hours of incubation. The fluorescence intensity was measured at 490/520 nm (excitation/emission) using a Microplate reader (Tecan, infinite 200 PRO, Switzerland).

### Statistics

Statistical analyses were performed with GraphPad PRISM software (GraphPad software, Inc.).

## Results

### Early BBB breakdown in prion-infected mice

The integrity of BBB was examined using C57Bl/6J mice inoculated intraperitoneally with mouse-adapted prion strain SSLOW [45, 46]. In mice, SSLOW causes profound, widespread neuroinflammation across multiple brain regions and induces more synchronous progression of neuropathology in different brain regions than other mouse-adapted strains [34, 38]. With the transformation of astrocytes into a reactive state, we observed a significant increase in AQP4 immunoreactivity within parenchymal astrocytic processes and a parallel decline in the localization of AQP4 with endfeet, which enwrap blood vessels (Fig. 1A,B,C). By the terminal stage of the disease (146 days post-inoculation or dpi), AQP4+ blood vessels were hardly noticeable, whereas the majority of AQP4 immunoreactivity was associated with parenchymal processes (Fig. 1A(*i*)). Remarkably, areas with aberrant localization of AQP4 were well noticeable already at the pre-symptomatic stage at 92 dpi and coincided with the areas of reactive astrogliosis (Fig. 1A(*ii*)). These observations revealed that astrocytes develop endfeet pathology in parallel with their transformation into a reactive state, which occurs at pre-symptomatic stages of the diseases.

**Figure 1.**
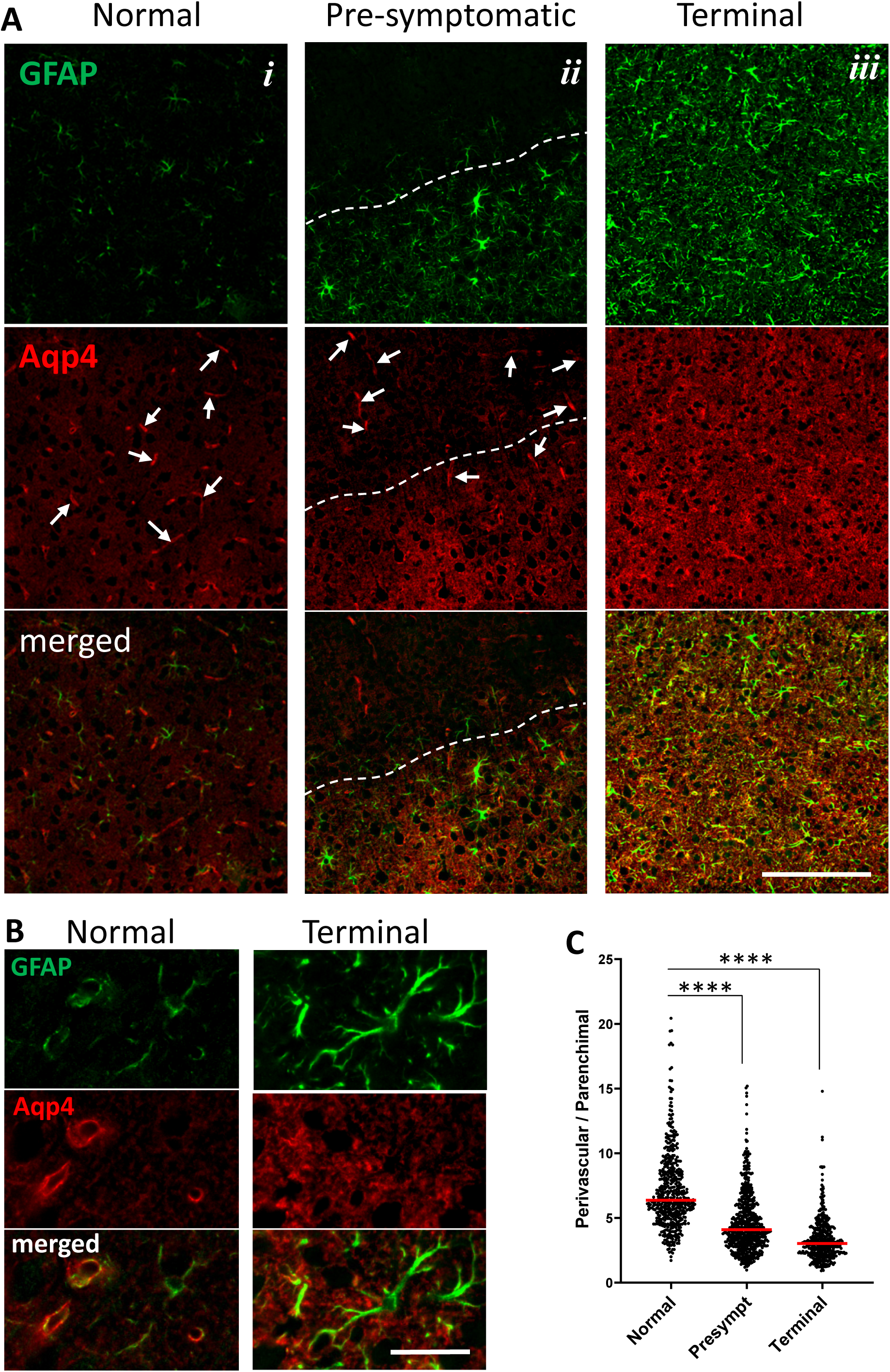
Aberrant localization of AQP4 in prion-infected mice. (**A**) GFAP and AQP4 immunofluorescence in the cortex of normal, age-matched mice (*i*), and SSLOW-infected mice at pre-symptomatic (92 dpi, *ii*) or terminal (146 dpi, *iii*) stages of the disease. Scale bar = 100 µm. Arrows point at AQP4+ blood vessels. White dashed line demark area with reactive astroglioses. (**B**) Magnified images of GFAP and AQP4 immunofluorescence in cortex of normal, age-matched mice (*left*) and terminally ill 146 dpi mice (*right*). Scale bar = 10 µm. (**C**) Ratio of perivascular to parenchymal AQP4 immunofluorescence in cortexes of normal, pre-symptomatic and terminal mice. Measurements were performed for 3 animals per group (3 fields of view for each brain, over 30 measurements from each field of view, resulting in a total of 493, 634 and 452 profiles for the normal, pre-symptomatic, and terminal mice, respectively. Red bars define mean. **** *p* < 0.0001 by Mann-Whitney test.

To assess BBB integrity, we employed Evans blue permeability assay, which involves injecting Evan blue into the mouse circulatory system and quantifying the dye in the brain parenchyma after perfusion. At the terminal stage of the disease, SSLOW-infected mice showed a significant increase in Evans blue fluorescence in brain parenchyma relative to that of age-matched control mice (Fig. 2A,B). Elevated levels of the dye were observed in multiple brain areas affected by prions and, in particular, the cortex and thalamus, two areas with the strongest neuroinflammation (Fig. 2A,B). No differences were found between age-matched mice, which were 6.5-7 months old, and 15-18 months old mice (referred to as aged group) (Fig. 2A-F), illustrating that the changes seen in prion-infected animals did not occur with normal aging. Examination of SSLOW-infected mice at the pre-symptomatic stage revealed elevated levels of Evans blue in the brain parenchyma, suggesting that a decline in BBB integrity begins prior to the clinical expression of the disease (Fig. 2A-F). Two methods for quantification of BBB permeability, one that examines the integrated intensity of Evans blue fluorescence of brain slices and another that relies on absorbance intensity of Evans blue extracted from brain regions, showed the same pattern of elevated Evans blue levels at pre-symptomatic and terminal stages relative to those in the age-matched control and aged animals (Fig. 2C-F). As a complementary approach for examining BBB integrity, the levels of IgGs in brain parenchyma were examined using Western blots. Again, SSLOW-infected animals showed considerably higher levels of heavy and light chain IgGs relative to the age-matched controls (Fig. 2G). In summary, these results documented a loss of BBB integrity in prion-infected mice, and showed that this process starts prior to the clinical expression of the disease.

**Figure 2.**
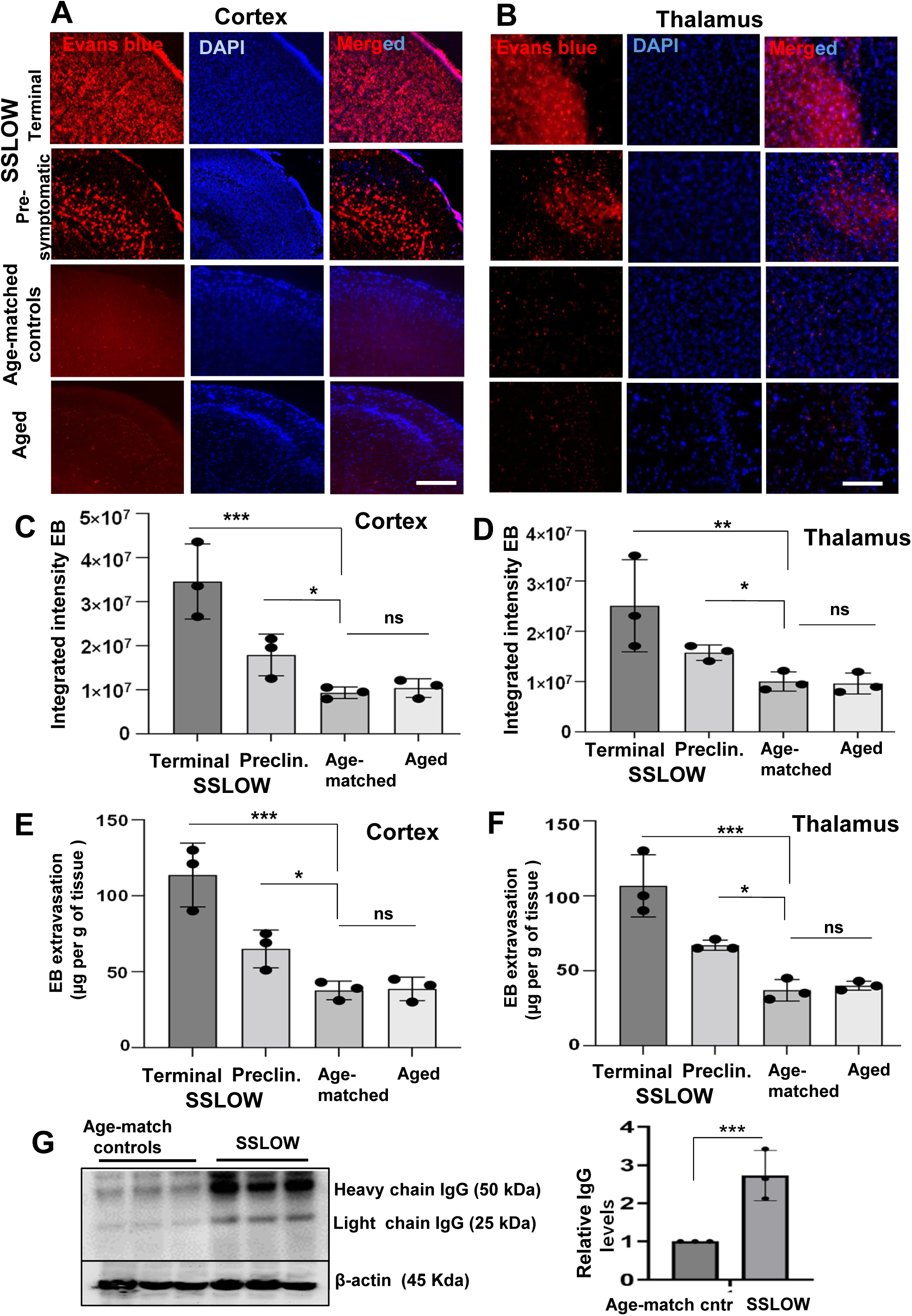
Loss of BBB integrity in prion-infected mice. **(A, B)** Fluorescence microscopy images of the cerebral cortex (**A**) and thalamus (**B**) from SSLOW-infected C57Bl/6J mice examined at the pre-symptomatic (110 dpi) and terminal (146-185 dpi) stages of the disease, aged-matched control (6.5-7 months old) and aged (15-18 months old) C57Bl/6J mice tested for BBB integrity using Evans blue permeability assay. Scale bars = 50 μm. (**C, D**) Quantification of integrated fluorescence intensity of Evans blue (EB) in the cerebral cortex (**C**) and thalamus (**D**) of four animal groups shown in panels **A** and **B**. (**E,F**) Quantification of Evans blue extracted from the cerebral cortex (**E**) and thalamus (**F**) in four animal groups shown in panels **A** and **B.** Quantification was based on measurements of Evans blue absorbance intensity. (**G**) Western blot and densitometric analysis of IgG heavy chain in cortical tissues of terminal SSLOW-infected mice and age-matched control mice. The IgG heavy chain signal is normalized per expression level of β-actin. In all experiments, mice were transcardially perfused. Data presented as the mean ± SE (*n*=3 animals per group), \*\*\**p*<0.001, \*\**p*<0.01, \**p*<0.05, ‘ns’ is non-significant by two-tailed, unpaired t-test. All groups are compared to the aged-matched control group.

### Brain endothelial junction proteins are downregulated in prion-infected mice

The integrity of the BBB is maintained by tight junctions between endothelial cells, which are complexes consisting of occludin and members of the claudin family, with Claudin-5 being the most abundant [47]. The *Zonula occludens* family members (ZO-1 and ZO-2) are localized in the cytoplasm and link the actin cytoskeleton with tight junctions via binding occludin. VE-cadherin (Cadherin 5) is a calcium-dependent transmembrane receptor and a member of the cadherin family that forms adherens junctions to provide stable adhesion between endothelial cells exclusively [48]. Unlike other cadherins, VE-cadherin is expressed only in endothelial cells. The most abundant component of junctions between endothelial cells is CD31 or PECAM-1, which does not directly participate in tight or adherens junctions, but helps maintain BBB integrity by forming trans-homophilic interactions between adjacent cells.

To get insight into the mechanism of BBB breakdown, we tested whether compromised BBB was associated with changes in the expression of tight junction and adherens junction proteins. RT-qPCR revealed downregulation of Occludin (*Ocln*), Claudin-5 (*Cldn5*) and VE-cadherin (*Cdh5*) in SSLOW-infected mice relative to age-matched controls, whereas the expression of ZO-1 (*TJP1*) and CD31 (*Pecam1*) did not change (Fig. 3A). Analysis of protein expression by Western blot confirmed downregulation of Occludin, Claudin-5 and VE-cadherin in SSLOW-infected brain tissues (Fig. 3 C, D, F), but no changes in the expression of ZO-1 or CD31 (Fig. 3B, E).

**Figure 3.**
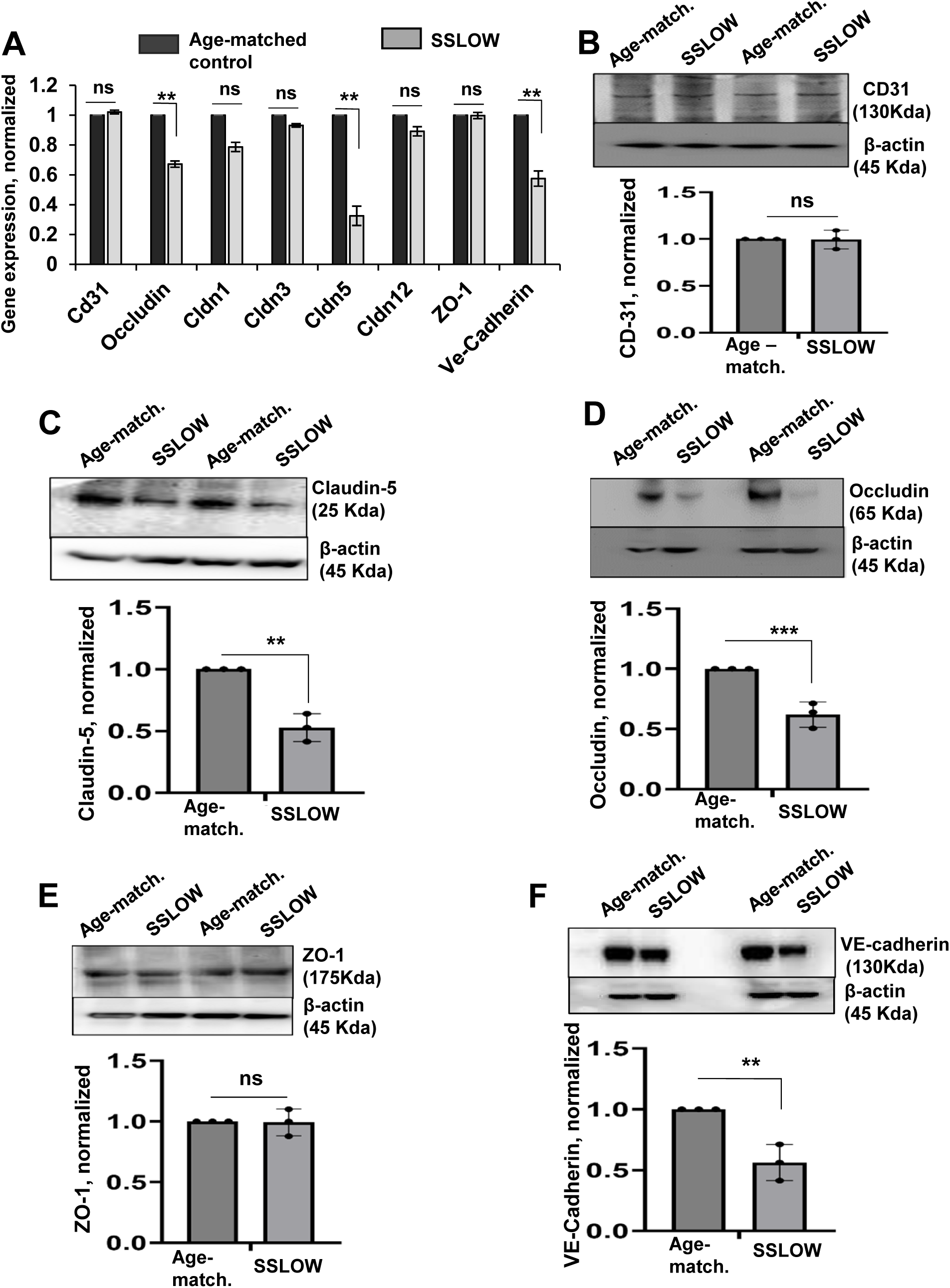
Brain endothelial junction proteins are downregulated in prion-infected mice. Expression of tight junction and adherens junction genes and proteins in SSLOW-infected C57Bl/6J mice at the terminal stage of the disease and age-matched control mice (6.5-7 months old) analyzed by RT-qPCR (**A**) or Western blot (**B-F**). (**A**) The gene expression in brain tissues of SSLOW-infected animals is normalized by the expression levels in age-matched control. *Gapdh* was used as a housekeeping gene. (**B-F)** Representative Western blots and quantitative analysis of CD-31 (**B**), Claudin-5 (**C**), Occludin (**D**), ZO-1 (**E**) and VE-cadherin (**F**) expression, which is normalized per expression of β-action, and plotted as the expression level in SSLOW-infected brains relative to the expression in aged-matched control mice. Data represent mean ± SE (*n*=3 animals per group), \*\*\**p*<0.001, \*\**p*<0.01 and \**p*<0.05 compared to aged-matched control or as indicated; ‘ns’ indicates non-significant, by two-tailed, unpaired t-test.

To investigate the impact of downregulation of endothelial junction proteins on vessel morphology, small blood vessels were analyzed using dual staining of brain slices for CD31 and one of the proteins associated with tight or adherens junctions, namely Occludin, Claudin-5, VE-cadherin or ZO-1 (Fig. 4). CD31 staining was used for the identification of CD31^+^ vasculature. Two parameters were analyzed: (i) integrated intensity of Occludin, Claudin-5, VE-cadherin or ZO-1 immunofluorescence, and (ii) length of Occludin^+^, Claudin-5^+^, VE-cadherin^+^ or ZO-1^+^ segments within CD31^+^ vasculature. In agreement with RT-qPCR and Western blot analyses, the signal intensity of Occludin, Claudin-5 and VE-cadherin within CD31^+^ vasculature was considerably lower in brains of SSLOW-infected mice relative to those of age-matched controls (Fig. 4A,B,D), while the intensity of the ZO-1 signal did not differ between the two groups (Fig. 4C). Staining for Occludin, Claudin-5 and VE-cadherin revealed segmented patterns within blood vessels in SSLOW-infected animals. In fact, the average length of Occludin^+^, Claudin-5^+^ or VE-cadherin^+^ segments was significantly shorter in SSLOW-infected mice relative to those of age-matched controls (Fig. 4A,B,D). Again, the average length of ZO-1^+^ vessels did not differ between SSLOW-infected and control groups (Fig. 4C). In summary, the above results suggest that in prion-infected mice, the breakdown of the BBB is associated with the downregulation of proteins that constitute tight and adherens junctions between endothelial cells. Consistent with a loss of BBB integrity, morphological analysis revealed gaps in tight and adherens junctions between endothelial cells of brain microvasculature.

**Figure 4.**
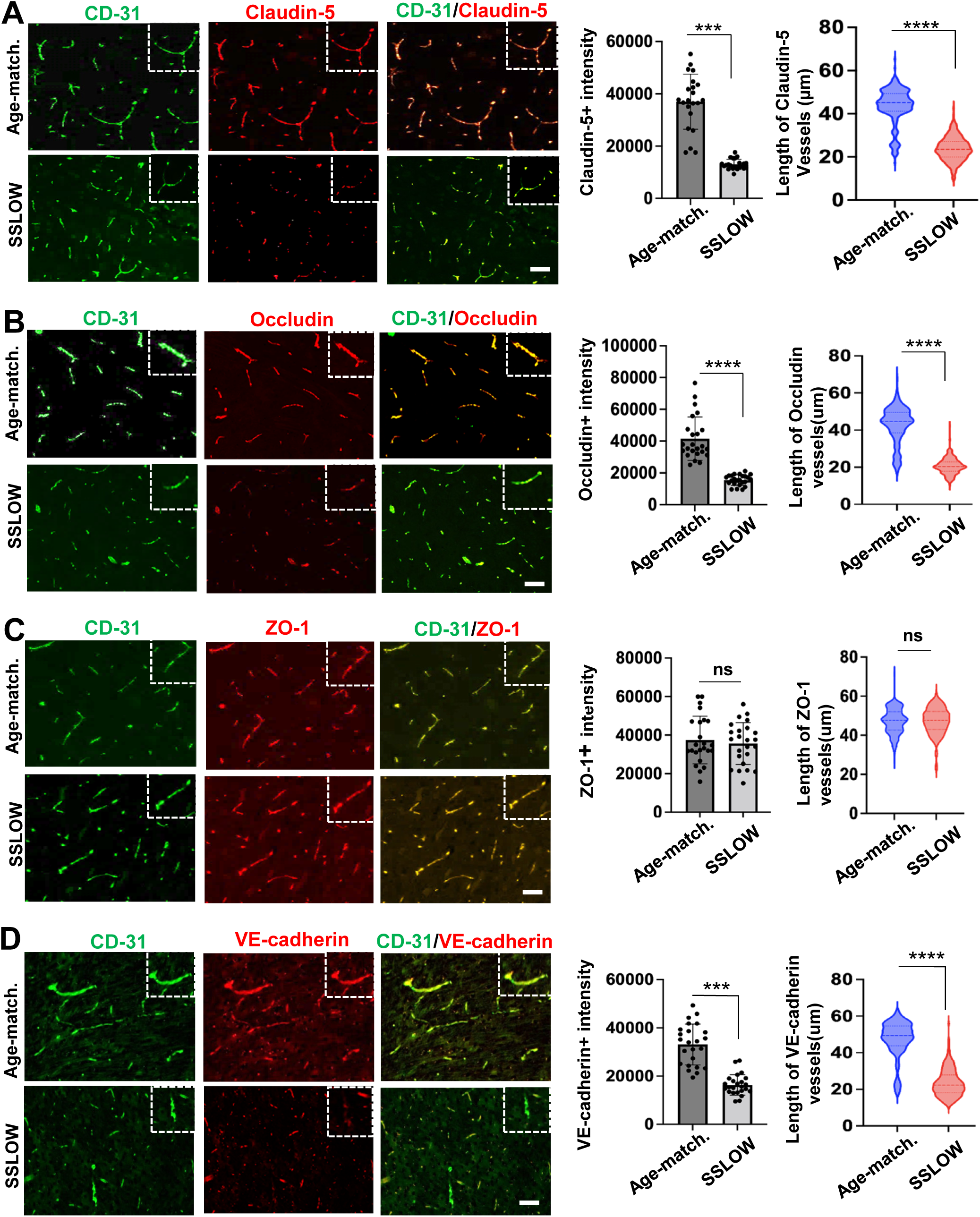
Analysis of blood vessel morphology in prion-infected brains. Representative immunofluorescence images of cerebral cortex from SSLOW-infected mice examined at the terminal stage of the disease and aged-matched control mice co-immunostained for endothelial capillary marker CD31 (green) and Claudin-5 (**A**, red), occludin (**B**, red), ZO-1 (**C**, red) or VE-cadherin (**D**, red). Right panels represent thee quantification of integrated fluorescence intensity and analysis of the length of claudin-5-, occludin-, ZO-1- or VE-cadherin - positive microvessels. For analysis of fluorescence intensity, the data represent mean ± SE, *n*=15 images from three animals in each group. For analysis of vessel length, *n*=400 vessels analyzed from three animals in each group. The dashed lines in violin plots show the median and quartiles. ****p<0.0001, ***p<0.001, ‘ns’ indicates non-significant, by two-tailed unpaired t-test with nonparametric Mann-Whitney test. Scale bars = 50 μm.

### Cell-to-cell junctions are impaired in primary endothelial cells that originate from prion-infected mice

To get insight into the mechanisms responsible for BBB breakdown, endothelial cells were isolated from clinically sick C57Bl/6J mice (142-185 dpi) that were infected intraperitoneally with SSLOW, or from age-matched control mice. The purity of the primary cultures was assessed through (i) co-immunostaining for the endothelial-specific marker CD31 along with the markers of astrocytes, microglia or oligodendrocyte (GFAP, Iba1 and OLIG2, respectively) (Fig. S2A), and (ii) RT-qPCR to analyze the expression of endothelial (*CD31*, *Ocln*, *Vwf*, *Glut1, Cldn1*, *Cldn5*, *ZO-1*, *Icam1)*, microglia (*Itgam*), neuron-(*Fox3*), and oligodendrocyte (*Mbp*) specific genes. When normalized relative to the expression levels in mouse cortex tissues, the endothelial primary cultures were enriched with the transcripts of endothelial specific genes, whereas the expression of markers of other cell types was found to be considerably low (Fig. S1B). Consistent with the gene expression analysis, co-immunostaining did not detect microglia, astrocytes or oligodendrocytes in primary endothelial cell cultures.

It has previously been demonstrated that endothelial cells isolated from mouse or human brains, when cultured *in vitro*, retain the ability to establish functional cell-to-cell tight and adherens junctions [49–51]. Upon analysis of brain endothelial cells (BECs) isolated from SSLOW-infected, clinically sick (SSLOW-BECs) and age-matched control mice (CT-BECs), significant differences were observed in cell morphology (Fig. 5A). CT-BECs exhibited continuous borders between neighboring cells positive for Occludin, Claudin-5 and VE-cadherin, indicating successful re-establishment of tight and adherens junctions in cultures (Fig. 5A,B). Conversely, in SSLOW-BECs, most cells lacked intercellular borders positive for Occludin, Claudin-5 or VE-cadherin (Fig. 5A,B), with these proteins displaying intracellular, often perinuclear localization (Fig. 5A). These findings suggest that SSLOW-BECs were unable to re-establish tight and adherens junctions, and instead maintained their disease-associated phenotype *in vitro*. Both SSLOW-BECs and CT-BECs expressed endothelial-specific genes (*Ocln*, *Cldn1*, *Cldn3*, *Cldn5*, *Cldn12, ZO-1*, *Vegfr2*, and *Cdh5*) (Fig. 5C). However, compared to CT-BECs, SSLOW-BECs exhibited significantly lower expression levels of genes (*Ocln*, *Cldn5*, *Cdh)* and corresponding proteins (Occludin, Claudin-5, VE-cadherin) responsible for tight and adherens junctions, as quantified by RT-qPCR and Western blot (Fig. 5C,D). Surprisingly, the expression of *Vegfr2* (Vascular Endothelial Growth Factor Receptor 2) was upregulated in SSLOW-BECs (Fig. 5C). Previous studies have shown that Vegfr2 mediates signaling by Vascular Endothelial Growth Factor A, which promotes BBB disruption [49, 52].

**Figure 5.**
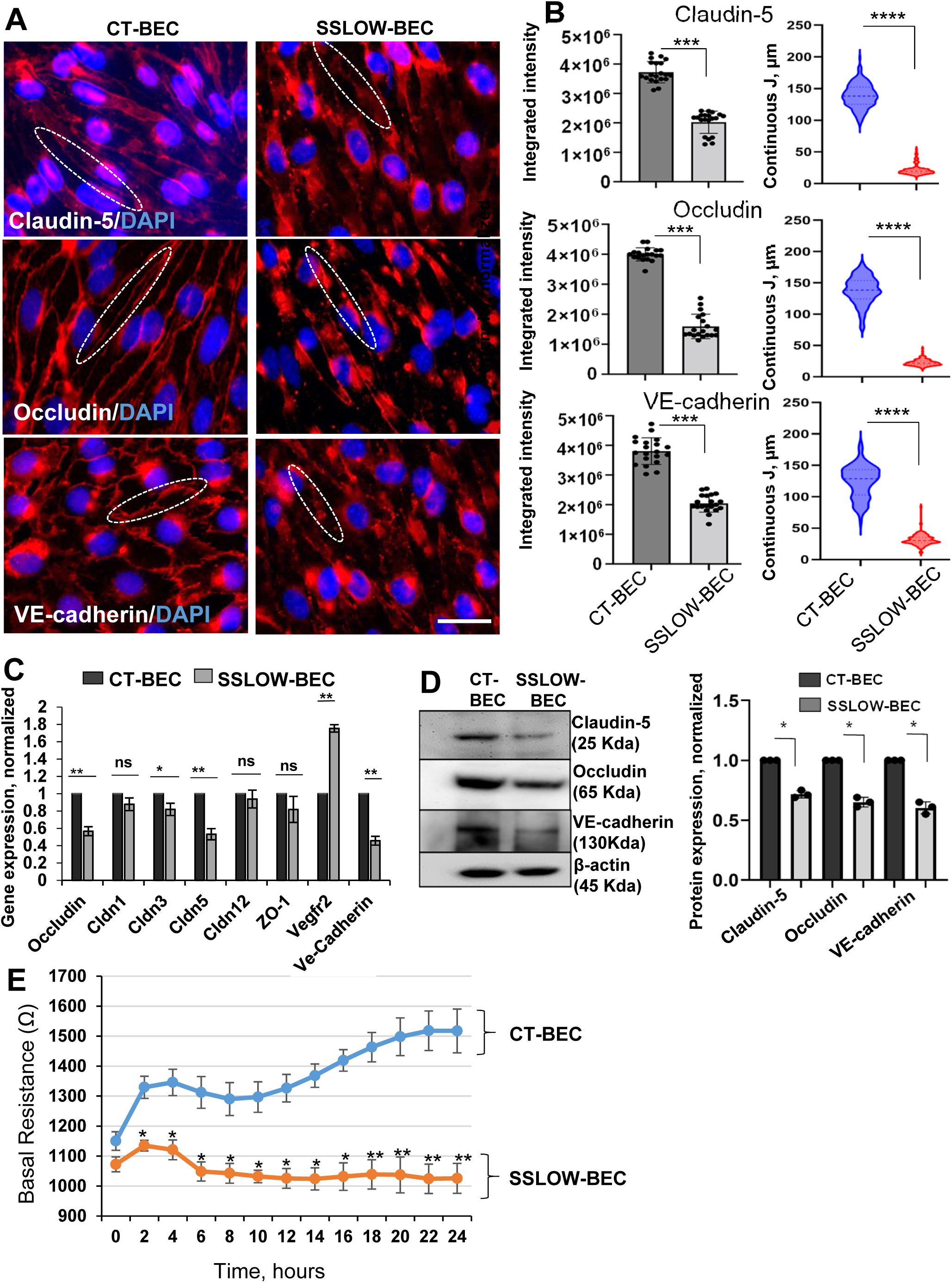
Endothelial cells isolated from prion-infected mice exhibit disease-associated phenotype. Primary endothelial cells (BECs) were isolated from SSLOW-infected mice (SSLOW-BECs) and aged-matched control mice (CT-BECs) and cultured for two to three weeks. (**A**) Immunofluorescence microscopy images of CT-BECs and SSLOW-BECs stained for Claudin-5, Occludin or VE-cadherin along with DAPI. Images are representatives of three cultures originating from independent animals. Dotted line encircles connected (CT-BECs) or disconnected (SSLOW-BECs) junctions between neighboring cells. Scale bars = 50 μm. (**B**) Quantification of integrated fluorescence intensity (left plots), and the length of discontinuous Claudin-5-, Occludin- or VE-cadherin-positive cell-to-cell junctions (right plots). For integrated intensity, *n*=20 random fields with 7–10 cells per field of view from three independent cultures, each prepared from an individual animal, per experimental group. For analysis of vessel length, *n*=150-200 discontinuous segments from three independent cultures, each prepared from an individual animal, per group. Data represent means ± SE, ****p<0.0001, ***p<0.001, by two-tailed unpaired t-test with nonparametric Mann-Whitney test. (**C**) Analysis of gene expression in SSLOW-BECs normalized by the expression in CT-BECs using qRT-PCR. *Gapdh* was used as a housekeeping gene. (**D**) Representative Western blots and densitometric analysis of Claudin-5, Occludin and VE-cadherin expression normalized per expression of β-action in CT-BECs and SSLOW-BECs. In **C** and **D**, data represent means ± SE, \*\**p*<0.01 and \**p*<0.05, ‘ns’ is non-significant by two-tailed, unpaired t-test. Data were collected for n=3 independent primary cell cultures per group, each prepared from an individual animal. (**E**) TEER assay of CT-BECs and SSLOW-BECs. Data were collected for *n*=3 independent primary cell cultures, each prepared from an individual animal, per group. For each independent culture, cells were plated and measured in triplicates. Data represent means ± SE, \*\**p*<0.01 and \**p*<0.05, by two-tailed, unpaired t-test.

To test whether changes in cell morphology and gene/protein expression impair physiological function, we employed electrical impedance sensing. This technique applies an alternating current between two electrodes separated by a cell monolayer to measure trans-endothelial electrical resistance (TEER). The cell monolayer acts as an insulator, dictating an impedance or resistance that quantifies the barrier properties of cultured cells. CT-BECs showed an increase in resistance over time, which is a sign of tighter cellular adhesion and the recovery of functional cell-to-cell junctions capable of maintaining barrier properties (Fig. 5E). In comparison to CT-BECs, SSLOW-BECs displayed a lower TEER, which also failed to increase with time (Fig. 5E). This result illustrates that the capacity for re-establishing of functional intercellular junctions is impaired in brain endothelial cells isolated from prion-infected animals.

### Reactive astrocytes from prion-infected animals impair the cell-to-cell junctions and barrier functions of healthy endothelial cells

To investigate whether reactive astrocytes are responsible for driving BBB dysfunction, primary astrocytes were isolated from SSLOW-infected, clinically sick mice (SSLOW-PACs) or age-matched control mice (CT-PACs) using a previously described method [6]. Compared to CT-PACs, SSLOW-PACs exhibited elevated levels of GFAP expression at both mRNA and protein levels (Fig. S2A). SSLOW-PACs also displayed an enlarged, hypertrophic morphology characterized by increased cell area, cell perimeter and a number of processes (Fig. S2B). Furthermore, when compared to CT-PACs, SSLOW-PACs showed upregulation of the expression of genes associated with astrocyte reactivity, along with pro-inflammatory genes known to be upregulated in prion-infected animals [24, 53] (Fig. S2C,D).

To co-culture primary endothelial cells with astrocytes, we utilized a Transwell membrane system that has been previously used as an *in vitro* model of BBB (Fig. 6A) [54]. After establishing SSLOW-PAC or CT-PAC cultures on the basolateral side of the Transwell membrane, CT-BECs isolated from non-infected adult animals were cultured as monolayers on the apical side of the membrane (Fig. 6A). In co-cultures with CT-PACs, endothelial cells formed long, uninterrupted cell-to-cell junctions positive for Occludin, Claudin-5 and VE-cadherin (Fig. 6B,C). However, in co-cultures with SSLOW-PACs, the lengths of Occludin-, Claudin-5- or VE-cadherin-positive segments between neighboring cells in CT-BECs were much shorter (Fig. 6B,C). In fact, CT-BECs co-cultured with SSLOW-PACs resembled SSLOW-BECs isolated from prion-infected animals, where Occludin, Claudin-5 and VE-cadherin localized predominantly in intracellular, often perinuclear sites (Fig. 5).

**Figure 6.**
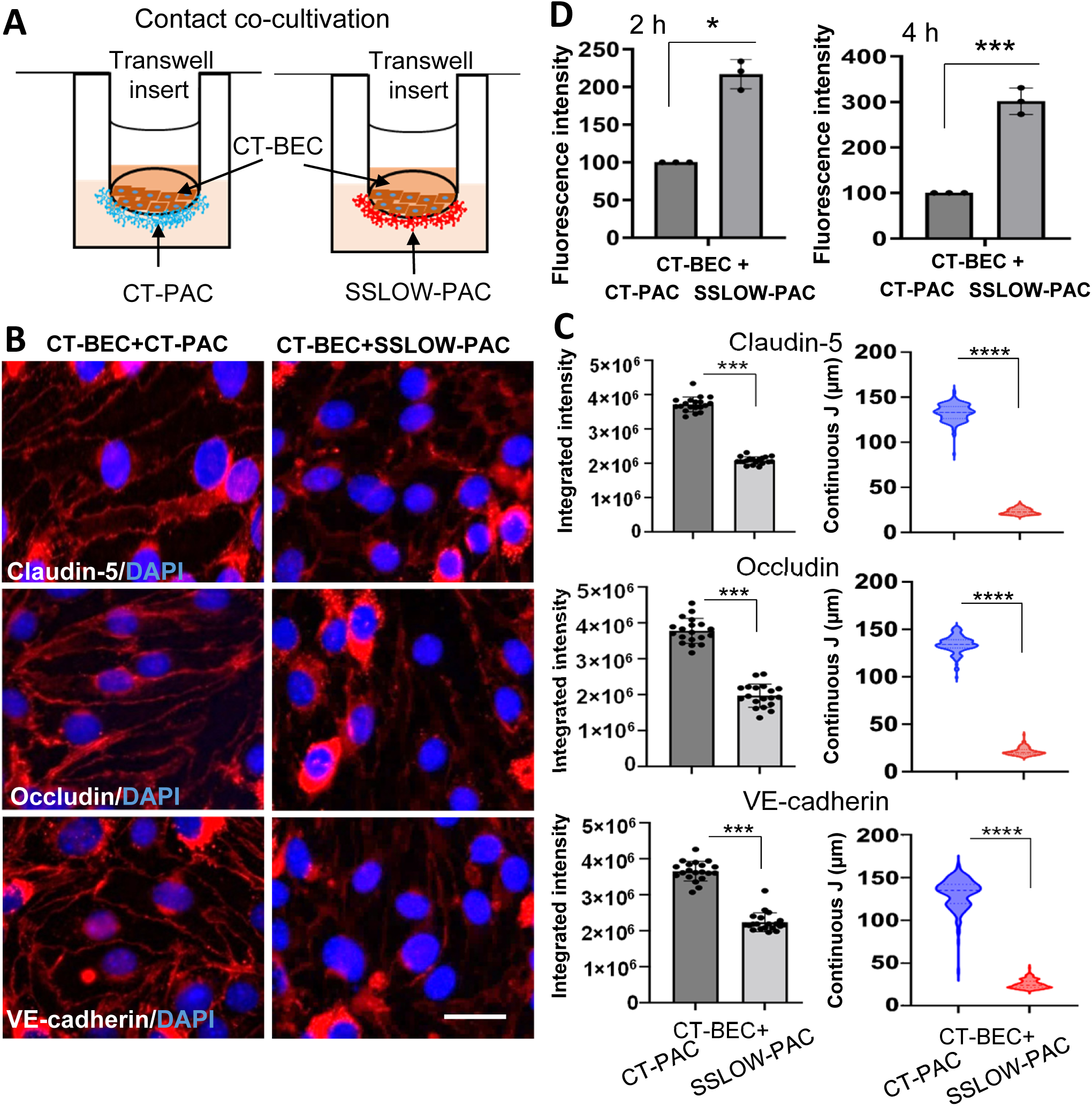
Reactive astrocytes from prion-infected animals impair cell-to-cell junctions and barrier functions of endothelial cells. (**A**) Schematic illustration of endothelial-astrocyte co-cultures in Transwell system. CT-BECs were grown on the apical side of inserts, whereas CT-PACs or SSLOW-PACs were cultured on the basolateral side of inserts. (**B**) Immunofluorescence microscopy images of CT-BECs co-cultured with CT-PACs or SSLOW-PACs and stained for Claudin-5, Occludin or VE-cadherin along with DAPI. Images are representatives of three cultures originating from independent animals. Scale bars = 50 μm. (**C**) Quantification of integrated fluorescence intensity (left plots), and the length of discontinuous Claudin-5-, Occludin- or VE-cadherin-positive cell-to-cell junctions (right plots). For integrated intensity, *n*=15 random fields with 5–10 cells per field of view from three independent co-cultures, each prepared from an individual animal, per experimental group. Data represent means ± SE, \*\*\**p*<0.001, by two-tailed unpaired t-test. For analysis of vessel length, *n*=150-200 continuous segments from three independent co-cultures, each prepared from an individual animal, per group. Data represent means ± SE, ****p<0.0001, by two tailed unpaired t test with nonparametric Mann-Whitney test. (**D**) Fluorescence intensity measured using FITC-dextran permeability assays in co-cultures of CT-BEC with CT-PAC or SSLOW-PAC after 2h or 4h of incubation with FITC-dextran. Data represent means ± SE of three independent co-cultures, each originating from individual animals. ***p<0.001 and *p<0.05, by two-tailed unpaired t-test.

To test the barrier properties of CT-BECs, we utilized the FITC-dextran permeability assay, which measures the diffusion of FITC-dextran through an endothelial cell monolayer in the Transwell system. We observed that CT-BECs cultured with SSLOW-PACs had significantly higher permeability compared to those co-cultured with CT-PACs (Fig. 6D). In conclusion, these findings demonstrate that reactive astrocytes from prion-infected animals hinder formation of cell-to-cell junctions and have adverse effects on the barrier properties of endothelial monolayers.

### Factors released by reactive astrocytes impair cell-to-cell junctions and barrier functions of endothelial cells

The negative effects of reactive astrocytes on cell-to-cell junctions raise the question of whether the deleterious effects are mediated by factors released by reactive astrocytes. Therefore, we examined the effect of media conditioned by reactive astrocytes isolated from prion-infected animals on cell-to-cell junctions and permeability of BECs isolated from normal, non-infected mice (Fig. 7A). Treatment of BECs with astrocyte-conditioned media form SSLOW-PACs (SSLOW-ACM) had a detrimental effect on the morphology and barrier functions of endothelial cells (Fig. 7B,C). The majority of Occludin, Claudin-5, or VE-cadherin immunoreactivity localized in intracellular perinuclear spaces, whereas only short stretches of Occludin-, Claudin-5-, or VE-cadherin-positive cell-to-cell junctions could be found between neighboring cells (Fig. 7B). This is in sharp contrast to BECs treated with astrocyte-conditioned media form CT-PACs (CT-ACM) that showed long, uninterrupted borders between cells positive for junction proteins (Fig. 7B). Upon treatment with SSLOW-ACM, TEER recorded for BECs declined with time and was considerably lower relative to TEER for non-treated BECs or BECs treated with CT-ACM (Fig. 7C). The time-dependent decline in TEER levels is indicative of cell retraction, loss of cell adhesion and cell-to-cell junctions. Assessment of cell viability revealed that a decline in TEER in SSLOW-ACM-treated BECs was not due to cell death, suggesting instead that the SSLOW-ACM impairs barrier properties of endothelial monolayers via altering cell morphology (Fig. 7B). In contrast to SSLOW-ACM, treatment of BECs with CT-ACM increased TEER in comparison to the resistance measured for non-treated BECs (Fig. 7C). As expected, BECs treated with CT-ACM maintained cell-to-cell junctions positive for Occludin, Claudin-5 and VE-cadherin (Fig. 7B). This result suggests that, in contrast to factors secreted by reactive astrocytes, factors released by normal astrocyte improve barrier characteristics of endothelial monolayers.

**Figure 7.**
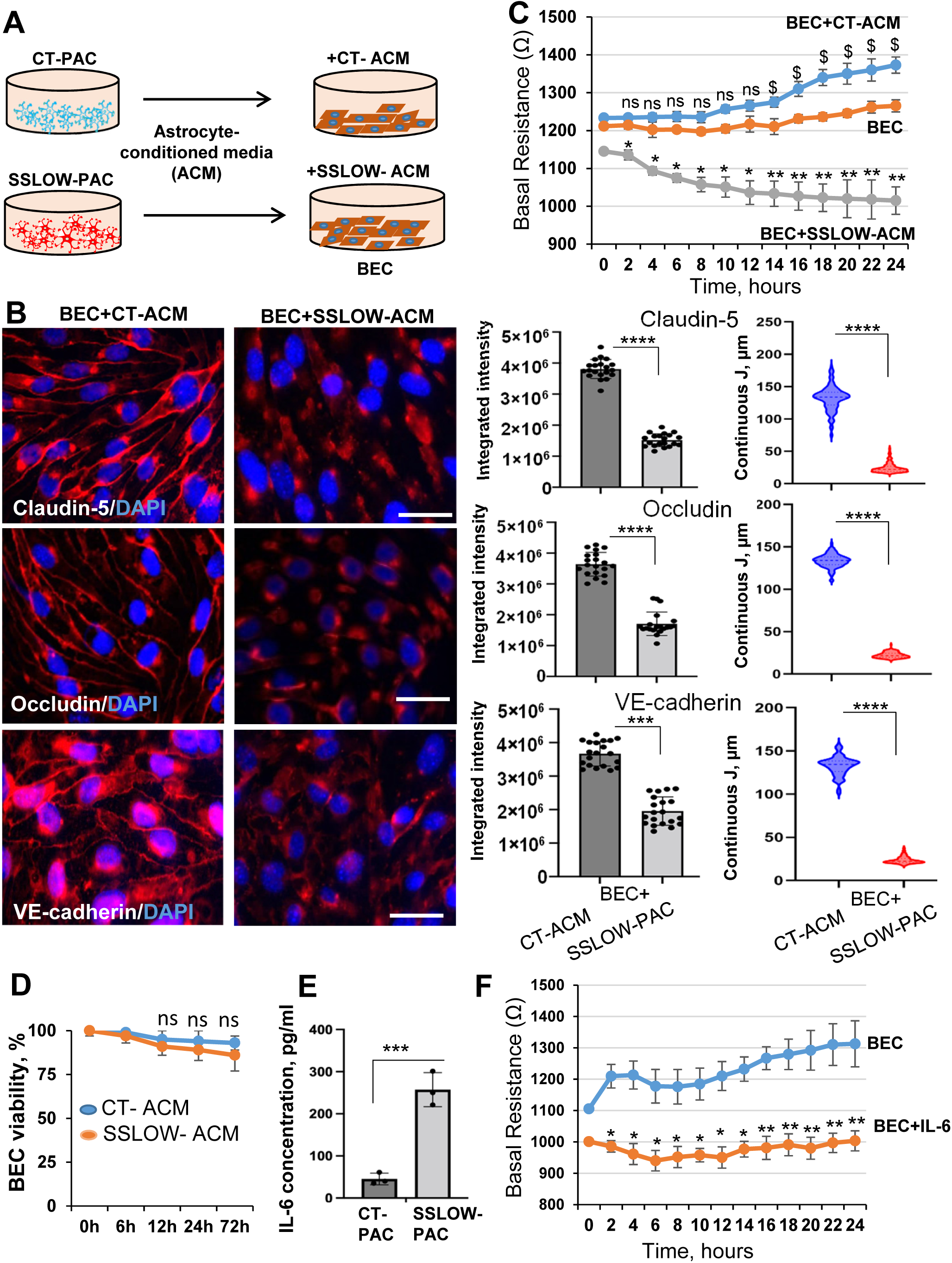
Factors released by reactive astrocytes impairs cell-to-cell junctions and barrier functions of endothelial cells. **(A)** Schematic diagram illustrating experimental design. (**B**) Panels on the left: immunofluorescence microscopy images of BECs treated with CT-ACM or SSLOW-ACM for 72 hours and stained for Claudin-5, Occludin or VE-cadherin along with DAPI. Images are representatives of three cultures originating from independent animals. Scale bar□=□50 µm.Panels on the right: Quantification of integrated fluorescence intensity (left plots), and the length of discontinuous Claudin-5-, Occludin- or VE-cadherin-positive cell-to-cell junctions (right plots). For integrated intensity, *n*=10 random fields with 7–10 cells per field of view from three independent co-cultures, each prepared from an individual animal, per experimental group. Data represent means ± SE, ****p<0.0001 and \*\*\**p*<0.001, by two-tailed unpaired t-test. For analysis of vessel length, *n*=150-200 continuous segments from three independent co-cultures, each prepared from an individual animal, per group. Data represent means ± SE, ****p<0.0001, by two-tailed unpaired t-test with nonparametric Mann-Whitney test. (**C**) TEER assay of non-treated BECs, and BECs pretreated with SSLOW-ACM or CT-ACM for 72 hours. Data were collected for *n*=3 independent primary cell cultures, each prepared from an individual animal, per group. For each independent culture, cells were plated and measured in triplicates. Data represent means ± SE, \*\**p*<0.01 and ^*, $^ *p*<0.05, ‘ns’ non-significant, by one-way ANOVA followed by Dunnett’s multiple comparison test, where BEC+CT-ACM and BEC+SSLOW-ACM were compared to BEC. (**D**) Cell viability in BECs assessed by MTT assay upon culturing in the presence of CT-ACM or SSLOW-ACM. The cell viability is expressed as a percentage relative to the viability of BECs treated with control-ACM at zero time point. (**E**) Analysis of IL-6 concentration in media conditioned by CT-PACs and SSLOW-PACs. In panels **D** and **E**, *n*=3 independent experiments representing three cultures originating from individual animals per group. For each independent culture, cells were plated and measured in triplicates. Data represent means□±□SE, \*\*\**p*<0.001, ‘ns’ non-significant, by two-tailed, unpaired t-test. **(F)** TEER assay of non-treated BECs, and BECs treated with IL-6 (10 ng/ml) for 72 hours. Data were collected for *n*=3 independent primary cell cultures, each prepared from an individual animal, per group. For each independent culture, cells were plated and measured in triplicates. Data represent means ± SE, \*\**p*<0.01 and \**p*<0.05, by two-tailed, unpaired t-test.

Among the factors secreted by reactive astrocytes is IL-6, which has been found to elevate BBB permeability and downregulate the expression of Claudin-5, Occludin and ZO-1 [55–58]. The secretion of IL-6 by astrocytes is known to be upregulated in prion diseases [6]. Consistent with previous studies, significantly higher levels of IL-6 were found in SSLOW-ACM relative to CT-ACM (Fig. 6E). Treatment with recombinant IL-6 decreased the TEER of BECs, supporting the idea that proinflammatory cytokines secreted by reactive astrocytes associated with prion diseases have detrimental effects on the barrier properties of brain endothelial cells (Fig. 7F). In summary, the above results demonstrate that the detrimental effects of reactive astrocytes on the morphology and function of endothelial cells are mediated via secreted factors.

### Astrocyte-derived extracellular vesicles reverse the disease phenotype of endothelial cells isolated from prion-infected animals

Previous experiments suggested that factors released by normal astrocytes can improve the barrier characteristics of endothelial cells isolated from adult, non-infected animals (Fig. 7C). This result is consistent with previous studies where extracellular vesicles (EVs) derived from immortalized astroglial cell line increased TEER and upregulated expression of tight junction proteins in a microvascular endothelial cell line [59]. Therefore, we decided to test whether EVs secreted by primary astrocytes isolated from adult, non-infected animals versus prion-infected animals can reverse the disease-associated phenotype of endothelial cells isolated from prion-infected animals. EVs were isolated from astrocytes originating from normal and prion-infected animals, and displayed EV markers Flotillin-1 and Alix (Fig. S3).

Consistent with previous results (Fig. 5A), SSLOW-BECs formed interrupted cell-to-cell junctions that were positive for Occludin, Claudin-5 or VE-cadherin, and all three proteins localized predominantly at intracellular, often perinuclear sites (Fig. 8A). As expected, TEER was considerably lower in SSLOW-BECs relative to CT-BECs (Fig. 8B). However, after treatment with EVs originating from normal astrocytes (CT-EVs), the TEER of SSLOW-BECs increased significantly, although it did not reach the TEER levels of CT-BECs (Fig. 8B). Consistent with the improvement in resistance, SSLOW-BECs treated with CT-EVs formed cell-to-cell junctions positive for Occludin, Claudin-5 or VE-cadherin (Fig. 8A). In CT-EVs-treated SSLOW-BECs, the length of uninterrupted cell-to-cell junctions increased significantly relative to those in non-treated SSLOW-BECs, although it did not reach the length of uninterrupted junctions seen in CT-BECs (Fig. 8A). Nevertheless, as judged from Western blot, treatment of SSLOW-BECs with CT-EVs upregulated the expression of all three proteins (Occludin, Claudin-5 and VE-cadherin) that are involved in cell-to-cell-junctions (Fig. 8C). Remarkably, treatment of CT-BECs with SSLOW-EVs downregulated the expression of Occludin, Claudin-5 and VE-cadherin (Fig. 8C). In summary, EVs from normal astrocytes upregulated the expression of cell junction proteins, improved TEER and partially reversed the disease phenotype of endothelial cells associated with prion-disease.

**Figure 8.**
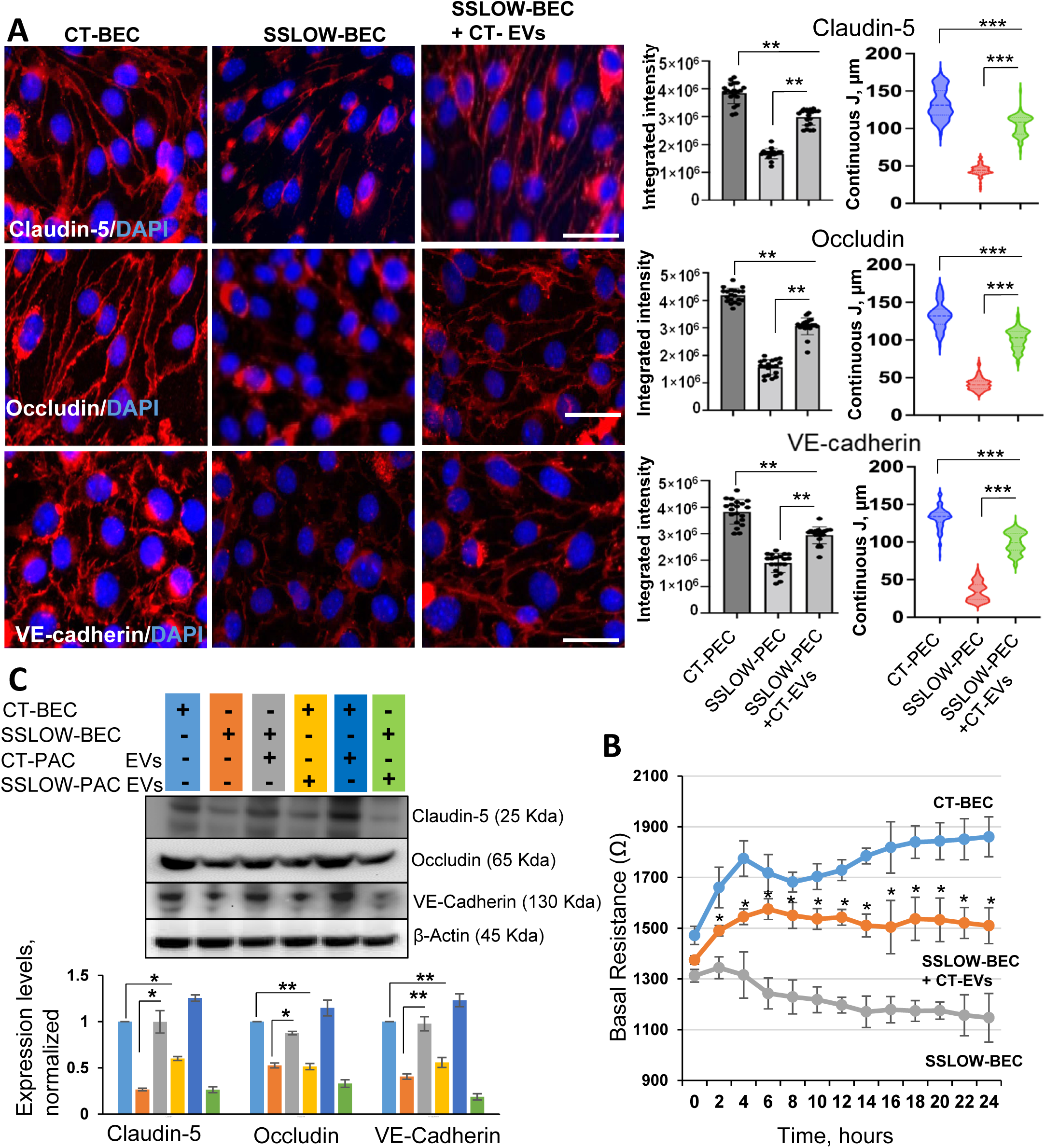
Astrocyte-derived EVs reverse the disease-associated phenotype of endothelial cells. (**A)** Panels on the left: immunofluorescence microscopy images of CT-BECs, SSLOW-BECs and SSLOW-BECs treated with CT-EVs (30µg/ml) for 72 hours and stained for Claudin-5, Occludin or VE-cadherin along with DAPI. Images are representatives of three cultures originating from independent animals. Panels on the right: Quantification of integrated fluorescence intensity (left plots), and the length of discontinuous Claudin-5-, Occludin- or VE-cadherin-positive cell-to-cell junctions (right plots). Scale bar□=□50 µm. For integrated intensity, *n*=20 random fields with 7– 10 cells per field of view from three independent co-cultures, each prepared from an individual animal, per experimental group. For analysis of vessel length, *n*=150-200 continuous segments from three independent co-cultures, each prepared from an individual animal, per group. Data represent means ± SE, ***p<0.001 and **p<0.01, by one-way ANOVA with Bonferroni multiple comparisons test. (**B**) TEER assay of CT-BECs, SSLOW-BECs and SSLOW-BECs treated with CT-EVs (30 µg/ml) for 72 hours. Data were collected for *n*=3 independent primary cell cultures, each prepared from an individual animal, per group. For each independent culture, cells were plated and measured in triplicates. Data represent means ± SE, \**p*<0.05, by two-tailed, unpaired t-test. **(C)** Representative Western blots and densitometric analysis of Claudin-5, Occludin and VE-cadherin expression normalized per expression of β-action in CT-BEC and SSLOW-BEC cultures in the absence of treatment or after treatment with SSLOW-PAC EVs or CT-PAC EVs for 72 hours. Data represent means□±□SE, n□=□3 independent cultures, each isolated from individual animals **p□<□0.01, *p□<□0.05, one-way ANOVA followed by Bonferroni multiple comparison test.

## Discussion

Astrocytes are essential for the development and maintenance of the BBB [60]. They communicate and support the BBB via enwrapping blood vessels with endfeet, and by secreting protective factors, including vascular endothelial growth factor (VEGF), fibroblast growth factor (FGF), glial-derived neurotrophic factor (GDNF), Apolipoprotein E (APOE), angiopoietins and others (reviewed in [8, 61]). However, the question of whether astrocytes continue to support the BBB in their reactive states, or become harmful to the BBB has been a subject of debate [61, 62].

To our knowledge, the current work is the first to document that, in prion disease, reactive astrocytes are detrimental to BBB integrity. A significant decline in the localization of AQP4 in astrocytic endfeet was observed at the pre-symptomatic stage of the disease, revealing that a loss of astrocyte polarity, along with perturbations in astrocyte-endothelial interactions, occurs early in the disease. In parallel with the loss of astrocyte polarity, Evans blue permeability assay demonstrated that a breakdown of the BBB occurred prior to the clinical expression of the disease. The loss in BBB integrity in prion-infected mouse brains was accompanied by a downregulation of key proteins that constitute tight and adherens junctions between endothelial cells including Occludin, Claudin-5 or VE-cadherin. Moreover, the morphological analysis of brain slices revealed gaps in tight and adherens junctions along blood vessels. Remarkably, in contrast to endothelial cells isolated from non-infected adult mice, the capacity of the endothelial cells originating from prion-infected animals for re-establishing functional intercellular junctions was impaired. In fact, substantially lower levels of Occludin, Claudin-5 and VE-cadherin, along with a decline in TEER, were observed in cells originating from prion-infected mice, relative to those of age-matched control mice. These results document that endothelial cells cultured *in vitro* preserved their disease-associated phenotype. Co-culture with reactive astrocytes from prion-infected animals or treatment with media conditioned by reactive astrocytes induced the disease phenotype in endothelial cells originating from non-infected adult mice. This phenotype involved downregulation and aberrant localization of Occludin, Claudin-5 and VE-cadherin, disruption of cell-to-cell junctions and an increase in permeability. Remarkably, treatment with EVs produced by normal astrocytes partially reversed the disease phenotype of endothelial cells isolated from prion-infected animals. Overall, our study provides experimental support to the hypothesis that reactive astrocytes drive pathological changes in endothelial cells, leading to BBB breakdown.

Astrocytes were found to respond to prion infection prior to neurons and even sooner than microglia [24]. Moreover, astrocyte functions scored at the top of the activated pathways [24, 63]. Transcriptome analysis of astrocyte-specific genes in prion-infected mice revealed a global perturbation across multiple homeostatic functions, including the loss of neuronal support function and changes in BBB support [34]. Consistent with transcriptome analysis, recent experiments that employed neuronal-astrocyte co-cultures revealed that the functions responsible for neuronal support, spine development, along with synapse maturation and integrity, were all impaired in reactive astrocytes isolated from the prion-infected mice [6]. Furthermore, a loss of important homeostatic functions was documented by a substantial decline in phagocytic activity in the reactive state associated with prion disease [64]. Interestingly, the degree of astrocyte reactivity was found to be predictive of the incubation time to prion disease, suggesting that phenotypic changes in astrocytes contribute to faster disease progression [34]. The current work demonstrates that, in addition to losses in homeostatic support of neuronal functions, reactive astrocytes are harmful to the integrity of the BBB.

AQP4, the most prevalent water channel in the CNS, is important for water homeostasis and the removal of neuronal waste from the brain via the glymphatic system [36, 65, 66]. AQP4 is localized in astrocyte endfeet that enwrap brain blood vessels. Through endfeet contacts, astrocytes regulate the expression of tight and adherens junction proteins, helping to maintain the integrity of the BBB [60]. In previous studies, tracking aberrant AQP4 localization, similar to that observed in the current work, was employed to report on the retraction of astrocytic endfeet from blood vessels in several neurological conditions, including major depressive disorder, ischemic insults, and vascular amyloidosis [67–71]. The aberrant localization of AQP4 in prion-infected brains suggests that the retraction of astrocytic endfeet from blood vessels also occurs in prion disease. We do not know whether the dissociation of astrocytic endfeet from vessels is one of the pathological events that trigger BBB dysfunction or, in contrast, a consequence of endothelial cell degeneration. Degeneration of endothelial cells was previously characterized by the downregulation and interrupted expression of tight junction proteins, a pathogenic signature often observed in neurodegenerative diseases [72, 73]. In fact, the dysfunction of tight and adherens junctions that leads to the loss of the polarity of endothelial cells and BBB breakdown has been emerging as a common feature among neurodegenerative diseases, including Alzheimer’s disease, Parkinson’s disease, Amyotrophic Lateral Sclerosis, multiple sclerosis and even normal aging [8, 74, 75]. In the current study, the downregulation of Occludin, Claudin-5 and VE-cadherin, along with the loss of BBB integrity, was observed in prion-infected mouse brains.

How do reactive astrocytes drive BBB breakdown? One potential mechanism is the downregulation of factors secreted by astrocytes that help to maintain BBB integrity. Alternatively, reactive astrocytes can drive BBB dysfunction via the upregulation of secreted inflammatory mediators. Astrocytes preserve region-specific identities in their reactive states and respond to prion infection in a region-specific manner [34, 76]. However, a common set of proinflammatory cytokines and other mediators contributing to neuroinflammation were found to be strongly upregulated irrespective of brain regions or prion strain [34]. Among these molecules are IL-6 and Serpina3n [6, 34]. Recent studies have shown that treatment with recombinant Serpina3n alone was sufficient to induce BBB dysfunctions in *ex vivo* mouse cortical explant cultures and in mice [77]. Similarly, members of the IL-6 family were previously found to downregulate the expression of Claudin-5, Occludin and ZO-1, and increase the permeability of the BBB [55–58, 78]. Consistent with the previous studies [6], upregulation of IL-6 secretion by reactive astrocytes originating from prion-infected mice was seen in the current work. Treatment of primary endothelial cells isolated from non-infected adult mice with recombinant IL-6 alone was sufficient to significantly reduce the integrity of monolayers as measured by TEER (Fig. 7E). While our work did not test pro-inflammatory cytokines in mixtures, it is likely that other molecules secreted by reactive astrocytes exacerbate the detrimental effect of IL-6 on BBB integrity even further. Indeed, the inflammatory stress induced by a mixture of IL-6, IL-17 and TNF-α was shown to be more detrimental to BBB integrity than the effects of individual cytokines [58]. The strong upregulation of IL-6 and Serpina3n in prion-infected mice suggests their plausible involvement in BBB breakdown.

Activation of Vascular Endothelial Growth Factor Receptors (VEGFR1 and VEGFR2) through VEGF-A during neurodevelopment leads to endothelial proliferation and differentiation. However, in the adult brain, the same pathway was found to promote BBB permeability [61]. Previous studies have shown that reactive astrocytes express VEGF-A and signal via VEGFR2 to promote BBB disruption in several neurological conditions [49, 52, 79, 80]. In CNS endothelial cultures, VEGF-A downregulates the expression of Claudin-5 and Occludin, while in a mouse brain, VEGF-A induces BBB breakdown leading to infiltration of immune cells [49, 52, 78, 81]. In the current study, upregulation of VEGFR2 was observed in endothelial cultures originating from prion-infected animals, suggesting that the VEGF-A/VEGFR2 signaling pathway might be involved in BBB breakdown in prion disease.

How does BBB dysfunction contribute to prion disease pathogenesis? BBB breakdown enables pathogens, peripheral immune cells and toxic blood-derived molecules to enter the brain in an uncontrolled manner. A significant increase of IgG in brain parenchyma shows that in prion-infected animals, the BBB is permeable to large proteins. The loss of BBB integrity is known to accelerate neuroinflammation, as peripheral cytokines and immune cells aggravate the activation of astrocytes and microglia (reviewed in [74]). Moreover, increased BBB permeability is associated with a decline in cerebral blood flow (reviewed in [82]). Although a loss of BBB integrity is likely to accelerate disease progression, it might also create an opportunity for pharmacological intervention via facilitating the delivery of drugs across the BBB as early as the onset of the disease.

Are pathological changes in the BBB reversible? Treatment with media conditioned by normal astrocytes improved the barrier properties of primary endothelial cells isolated from non-infected adult mice (Fig. 7C), supporting the view that astrocytes play a role in maintaining BBB integrity. Remarkably, treatment of endothelial cells isolated from prion-infected mice with EVs produced by normal astrocytes partially reversed the disease phenotype and improved their TEER (Fig. 8). These results raise the possibility that disease-associated changes in the BBB could be reversed *in vivo* by reversing astrocyte reactivity and restoring their homeostatic sate.

The current study does not address the question of whether reactive microglia also drive BBB dysfunctions. Previously, activation of microglia was shown to trigger BBB dysfunctions via proinflammatory cytokines (reviewed in [74]). A number of proinflammatory cytokines including Il1β, Oncostatin M and TNF-α that were found to induce BBB dysfunctions are strongly upregulated in prion disease too [24, 83]. It is likely that in addition to reactive astrocytes, reactive microglia contribute to the proinflammatory environment harmful to the BBB. It would be interesting to explore next whether targeting the BBB and/or restoring the homeostatic state of astrocytes will create opportunities for slowing down or reversing the progression of prion diseases.

## Conclusions

In prion diseases, astrocyte endfeet pathology, along with a progressive loss in BBB integrity, develops at the pre-symptomatic stage of the disease. Reactive astrocytes or media conditioned by reactive astrocytes, isolated from prion-infected mice, induced a disease-associated phenotype in endothelial cells originating from normal adult mice. Conversely, EVs produced by normal astrocytes partially reversed the disease-associated phenotype of endothelial cells isolated from prion-infected mice. This study demonstrate that, in prion disease, reactive astrocytes drive pathological changes in BBB. Moreover, our findings illustrate that the harmful effects are linked to proinflammatory factors secreted by reactive astrocytes.

## Abbreviations

ACM: Astrocyte conditioned media
SSLOW-ACM: Media conditioned by reactive astrocytes originating from SSLOW-inoculated mice
CT-ACM: Media conditioned by normal astrocytes originating from age-matched control mice
AQP4: Aquaporin 4
BBB: Blood brain barrier
BECs: Brain endothelial cells
SSLOW-BECs: BECs isolated from SSLOW-infected clinically sick mice
CT-BECs: BECs isolated from age-matched control mice
CD31: Cluster of differentiation also known as PECAM-1
CNS: Central nervous system
Dpi: Days post-inoculation
EVs: Extracellular vesicles
FITC: Fluorescein isothiocyanate
GFAP: Glial fibrillary acidic protein
IL: Interleukin
PACs: Primary astrocyte cells
SSLOW-PACs: PACs isolated from SSLOW-infected clinically sick mice
CT-PACs: PACs isolated from age-matched control mice
PECAM-1: Platelet and endothelial cell adhesion molecule 1 also known as CD31
PrP^C^: Normal cellular form of the prion protein
PrP^Sc^: Disease-associated infectious form of the prion protein
RT-qPCR: Reverse transcription-quantitative polymerase chain reaction
SSLOW: Mouse-adapted prion strain of synthetic origin
TEER: Trans-endothelial electrical resistance
TNF: Tumor Necrosis Factor
VEGF: Activation of Vascular Endothelial Growth Factor
VEGFR: Activation of Vascular Endothelial Growth Factor Receptor
ZO-1: Zonula occludens-1

## Declarations

### Ethics approval and consent to participate

The study was carried out in strict accordance with the recommendations in the Guide for the Care and Use of Laboratory Animals of the National Institutes of Health. The animal protocol was approved by the Institutional Animal Care and Use Committee of the University of Maryland, Baltimore (Assurance Number: A32000-01).

### Consent for publication

Not applicable

### Availability of data and materials

All data generated or analyzed during this study are included in this published article and its supplementary information file.

### Authors’ contributions

IB and RK designed the study; RK and KM performed animal procedures; RK performed experiments; RK analyzed the data; NM and NP analyzed AQP4 localization; YL and KB assisted with TEER measurements and TEER data analysis; IB and RK wrote the manuscript. All authors approved the final manuscript.

## Acknowledgments

Not applicable

## Supplemental figure legends

**Fig. S1. Isolation of adult endothelial cell cultures. (A)** Representative fluorescent microscopy images of primary endothelial cells isolated from adult C57BL/6J mice co-immunostained for CD31 (endothelial marker) and GFAP (astrocytes marker), Iba1 (microglia marker), or olig2 (oligodendrocytes marker). Nuclei are stained with DAPI. Scale bar = 50 μm. (**B)** Analysis of gene expression using qRT-PCR in BECs, normalized by the expression levels in cortical brain tissues. Gapdh was used as a housekeeping gene. Data represent means ± SE, *n*=3 independent cultures isolated from individual animals, \*\*\**p*<0.001, \*\**p*<0.01 and \**p*<0.05, by two-tailed, unpaired t-test.

**Fig. S2. Primary astrocytes isolated from prion-infected preserve reactive phenotype.** Primary astrocytes were isolated from SSLOW-infected, clinically ill mice at 142-185 dpi and non-infected, age-matched control mice and cultured in humidified conditions for two-to-three weeks. (**A**) Representative Western blots and densitometric analysis of GFAP expression normalized per expression of β-actin in CT-PACs and SSLOW-PACs. (**B)** Representative images of CT-PACs and SSLOW-PACs stained for GFAP, and morphometric analyses of cell area, perimeter and process number in astrocytes from CT-PACs and SSLOW-PACs. Insets show magnified images. Images are representatives of three independent cultures. Scale bar = 50 μm. **(C)** Analysis of expression of genes associated with astrocyte reactivity in SSLOW-PACs normalized by the expression levels in CT-PACs using qRT-PCR. (**D)** Analysis of expression of pro-inflammatory genes in SSLOW-PACs normalized by the expression levels in CT-PACs using qRT-PCR. Data represent means ± SE, *n*=3 independent primary cell cultures, each prepared from an individual animal, per group. \*\*\**p*<0.001, \*\**p*<0.01 and \**p*<0.05, ‘ns’ indicates non-significant, by two-tailed unpaired t-test.

**Fig. S3. Preparation of EVs from CT-ACM and SSLOW-ACM**. Representative Western blots of flotillin-1 (**A**) and Alix (**B**) in EVs isolated from CT-ACM and SSLOW-ACM. Data show three independent preparations of EVs from primary astrocyte cell cultures, each isolated from an individual animal.

