## Supplementary material for "Reactive astrocytes associated with prion disease impair the blood brain barrier": Table S1

**Table S1**. Primer sequences for RT-qPCR

| Primer | Accession number | Sequence |
| --- | --- | --- |
| Pecam1 | NM_008816.3 | F 5’- GAGAGTCCTGTGCACGTATTT-3’ |
|  |  | R 5’- GTTGTTGTAGCCAGCCATTTC-3’ |
| Cldn1 | NM_016674.4 | F 5’- TTTCAGGTCTGGCGACATTAG-3’ |
|  |  | R 5’- AGGTGTTGGCTTGGGATAAG-3’ |
| Cldn3 | NM_009902.4 | F 5’- CACCCACCAAGATCCTCTATTC-3’ |
|  |  | R 5’- GCGAGGTTTCTTTGTCCATTC-3’ |
| Cldn5 | NM_013805.4 | F 5’- CGGGTGAGCATTCAGTCTTTA-3’ |
|  |  | R 5’- CCGCCCTTAGACATAGTTCTTC-3’ |
| Cldn12 | NM_001193659.1 | F 5’- TGTCCTTCCTGTGTGGTATTG-3’ |
|  |  | R 5’-CAGCAGGAGAGCAGTCATAAA-3’ |
| Zo-1 | NM_001198985.2 | F 5’-CGAAACTGATGCTGTGGATAGA-3’ |
|  |  | R 5’-GAATGGCTCCTTGTGGGATAA-3’ |
| Cdh5 | NM_009868.4 | F 5’-CATCTAGGGTTCTGGTCTTTGG-3’ |
|  |  | R 5’-CTTTCTGGTGAGTGGGTTAGAG-3’ |
| Vegfr2 | NM_001363216.1 | F 5’- ATCCAGATGAGGGCAAGTTTAG-3’ |
|  |  | R 5’- TAATGAGAGCCACGCCATTAG-3’ |
| Vwf | NM_011708.4 | F 5’-GACCCAGTGCTGTGATGAATA-3’ |
|  |  | R 5’-GCCTTCGTGGAGAACATAAGA-3’ |
| Ocln | NM_001360536.1 | F 5’-CAAGAGGATGGTGGGAGATTATG-3’ |
|  |  | R 5’-GGCTGCTGCAAAGATTGATTAG-3’ |
| Glut1 | NM_011400.3 | F 5’- CCTCGTGCTCTTCTTCATCTT-3’ |
|  |  | R 5’-GGAGATAGGAGAGTGGCTGATA-3’ |
| Icam1 | NM_010493.3 | F 5’-CCAGTAGACACAAGCAAGAAGA-3’ |
|  |  | R 5’-GCATCCTGACCAGTAGAGAAAC-3’ |
| Itgam | NM_001082960.1 | F 5′- CAGCCTTTGACCTTATGTCATGG-3’ |
|  |  | R 5′- CCTGTGCTGTAGTCGCACT-3’ |
| Fox3 | NM_001039167.1 | F 5′-ACCTACAGCATCGGAACCAT- 3’ |
|  |  | R 5′-TTGCTAGTAGGGGGTGAAGC-3’ |
| Mbp | NM_001025251.2 | F 5′- CTCAGAGGACAGTGATGTGTTT-3’ |
|  |  | R 5′- CGCCTTGCCAGTTATTCTTTG-3’ |
| Lcn2 | NM_008491.1 | F 5′- CCCCATCTCTGCTCACTGTC–3’ |
|  |  | R 5′- TTTTTCTGGACCGCATTG-3’ |
| Serpina3n | NM_009252.2 | F 5′- GCAACACCCTGGAAGAGATT-3’ |
|  |  | R 5′- CTGGGTCTGTTTCCTCACATAG-3’ |
| Steap4 | NM_054098.3 | F 5′- CCCGAATCGTGTCTTTCCTATAA-3’ |
|  |  | R 5′- CCTCGATAGAGCTGCAGAATG-3’ |
| Cxcl10 | NM_021274.2 | F 5′- AGTAACTGCCGAAGCAAGAA-3’ |
|  |  | R 5′- GCACCTCCACATAGCTTACA-3’ |
| Timp1 | NM_001044384.1 | F 5′- CCTGGGTTTTCATTGGAGGGTTG-3’ |
|  |  | R 5′- GTGGCAGAACTTGAGGAGGTC-3’ |
| Serping1 | NM_009776.3 | F 5′- TGATGGCGCCTTTCTTCTAC-3’ |
|  |  | R 5′- CCACCTTGGCCTTCAAAGTA-3’ |
| H2t23 | NM_010398.3 | F 5′- GATACCTACGGCTGGGAAATG-3’ |
|  |  | R 5′- GTGATGTCAGCAGGGTAGAAG-3 |
| Ggta1 | NM_001308300.1 | F 5′- TCATCGGGTCCTACCTACAA-3’ |
|  |  | R 5′- CTTCAGTCACCTGCTCCATAC-3’ |
| S100a10 | NM_009112.2 | F 5′- GGGCTTCCAGAGCTTTCTATC-3’ |
|  |  | R 5′- CTCCAGTTGGCCTACTTCTTC-3’ |
| Tgm1 | NM_001161715.1 | F 5′- GCCCTTGAGCTCCTCATTG-3’ |
|  |  | R 5′- CCCTTACCCACTGGGATGAT-3’ |
| Il6 | NM_031168.2 | F 5′- TGCTGCCACCTTATTTACTCC-3’ |
|  |  | R 5′- AGTGGGGCATGATGACAGTT-3’ |
| Il33 | NM_001164724.2 | F 5′- TCCACGGGATTCTAGGAAGA-3’ |
|  |  | R 5′- GAGGCAGGAGACTGTGTTAAA-3’ |
| Il12b | NM_001303244.1 | F 5′- GCTTCTTCATCAGGGACATCA-3’ |
|  |  | R 5′- CTTGAGGGAGAAGTAGGAATG-3’ |
| Ccl2 | NM_011333.3 | F 5′- TGAGGCTCAGCACAGCAA-3’ |
|  |  | R 5′- ATGGGCTTCCAGAATACCG-3’ |
| Ccl4 | NM_013652.2 | F 5′- GCCCTCTCTCTCCTCTTGCT-3’ |
|  |  | R 5′- GGAGGGTCAGAGCCCATT-3’ |
| Ccl5 | NM_013653.3 | F 5′- TGCAGAGGACTCTGAGACAGC-3’ |
|  |  | R 5′- GAGTGGTGTCCGAGCCATA-3’ |
| Gapdh | NM_001289726.1 | F 5′- AACAGCAACTCCCACTCTTC-3’ |
|  |  | R 5′- CCTGTTGCTGTAGCCGTATT-3’ |
