## Supplementary figures and images for "Reactive astrocytes associated with prion disease impair the blood brain barrier"

### Figure S1

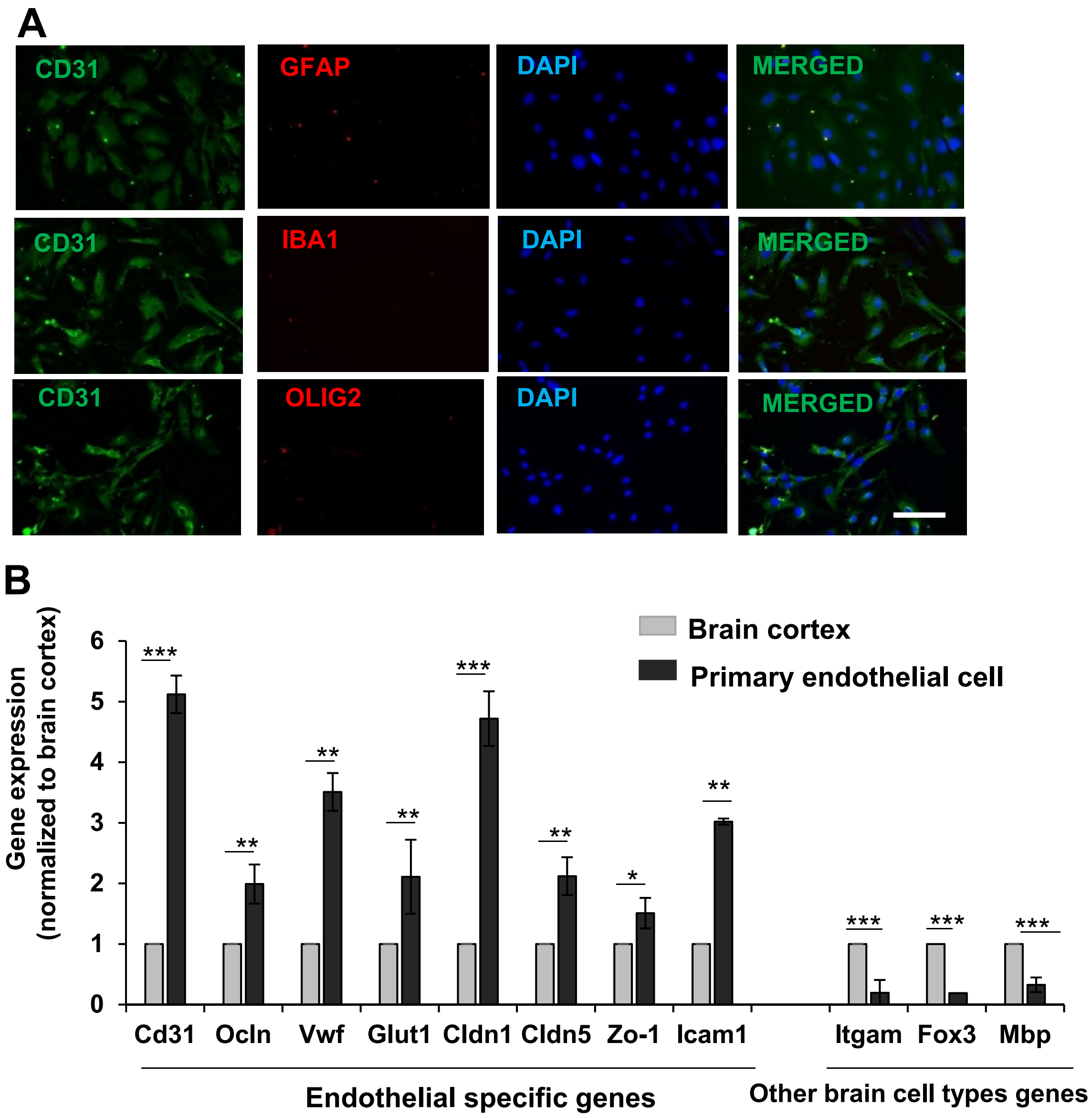

### Figure S2

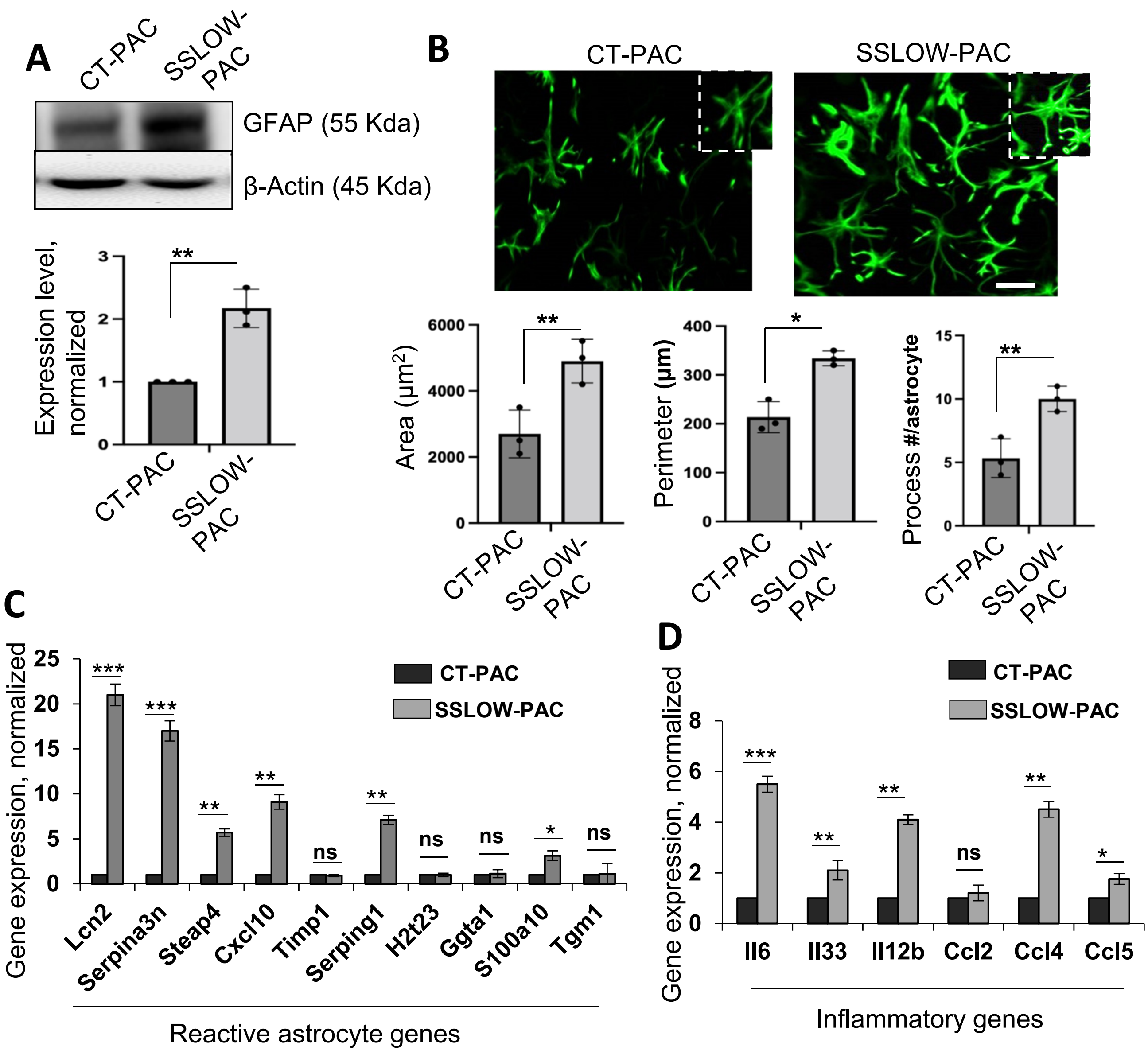

### Figure S3

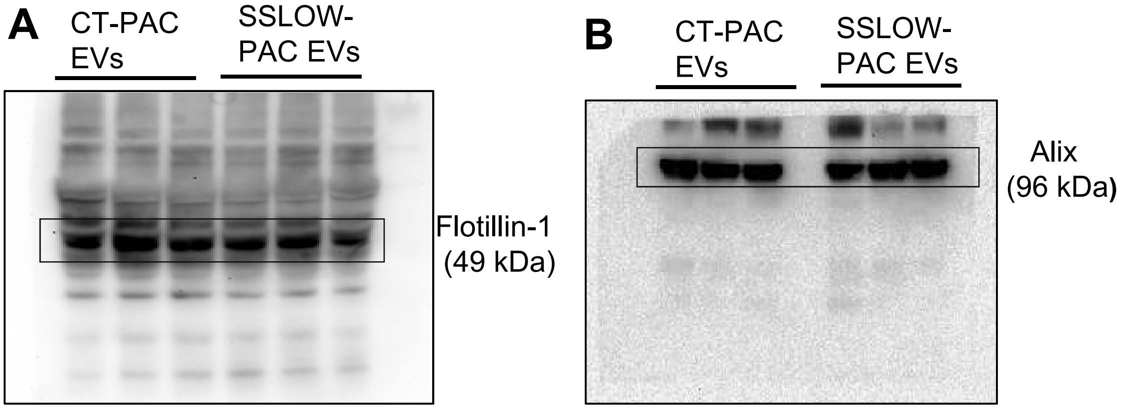
